# The evolution of social parasitism in *Formica* ants revealed by a global phylogeny

**DOI:** 10.1101/2020.12.17.423324

**Authors:** Marek L. Borowiec, Stefan P. Cover, Christian Rabeling

**Author notes:** Corresponding author. *Email address* (M. L. Borowiec). Corresponding author. *Email address* (C. Rabeling).

## Abstract

Studying the behavioral and life history transitions from a cooperative, eusocial life history to exploitative social parasitism allows for deciphering the conditions under which changes in behavior and social organization lead to diversification. The Holarctic ant genus *Formica* is ideally suited for studying the evolution of social parasitism because half of its 176 species are confirmed or suspected social parasites, which includes all three major classes of social parasitism known in ants. However, the life-history transitions associated with the evolution of social parasitism in this genus are largely unexplored. To test competing hypotheses regarding the origins and evolution of social parasitism, we reconstructed the first global phylogeny of *Formica* ants and representative formicine outgroups. The genus *Formica* originated in the Old World during the Oligocene (∼30 Ma ago) and dispersed multiple times to the New World. Within *Formica*, the capacity for dependent colony foundation and temporary social parasitism arose once from a facultatively polygynous, independently colony founding ancestor. Within this parasitic clade, dulotic social parasitism evolved once from a facultatively temporary parasitic ancestor that likely practiced colony budding frequently. Permanent social parasitism evolved twice from temporary social parasitic ancestors that rarely practiced colony budding, demonstrating that obligate social parasitism can originate from different facultative parasitic backgrounds in socially polymorphic organisms. In contrast to inquiline ant species in other genera, the high social parasite diversity in *Formica* likely originated via allopatric speciation, highlighting the diversity of convergent evolutionary trajectories resulting in nearly identical parasitic life history syndromes.

## Introduction

The complex societies of eusocial insects are vulnerable to exploitation by social parasites that depend on their host colonies for survival and reproduction without contributing to colony maintenance and brood care (Hölldobler and Wilson, 1990; Schmid-Hempel, 1998; Buschinger, 2009; Rabeling, 2020). Social parasitism is common among eusocial Hymenoptera and evolved independently in distantly related lineages, including bees, wasps, and ants (Wcislo, 1987; Cervo, 2006; Buschinger, 2009; Smith et al., 2013; Miller et al., 2015; Lhomme and Hines, 2019; Rabeling, 2020). Many studies on social parasitism have focused on the evolution of cooperation and conflict in colonies of eusocial insects and on co-evolutionary arms race dynamics between hosts and parasites (Foitzik et al., 2001; Brandt et al., 2005; Kilner and Langmore, 2011; Feldmeyer et al., 2017). However, the evolutionary origins of social parasitism and the co-evolutionary factors causing speciation and thereby contributing to the high diversity of social parasite species in eusocial insects are not well understood (Bourke and Franks, 1991; Huang and Dornhaus, 2008). Comparative evolutionary studies of social parasites are promising, because they are expected to provide novel insights into the conditions associated with a behavioral change from eusociality to social parasitism as well as into the consequences the life history transitions have on speciation and biological diversification.

Social parasitism is a life history strategy that evolved at least 60 times in ants, and more than 400 socially parasitic species are known from six distantly related subfamilies (Rabeling, 2020). Despite the high diversity, three main life history strategies can be recognized across social parasites: (i) temporary, (ii) dulotic, and (iii) inquiline social parasitism (Wasmann, 1891; Wheeler, 1904, 1910; Wilson, 1971; Hölldobler and Wilson, 1990; Kutter, 1968; Buschinger, 1986, 1990, 2009). The queens of temporary socially parasitic ant species invade the host nest, kill the resident queen(s), and the host workers raise the parasite’s offspring (Wheeler, 1904). In the absence of an egg-laying host queen, the host workforce is gradually replaced until the colony is composed solely of the temporary social parasite species. The queens of dulotic social parasites start their colony life cycle as temporary social parasites, and once sufficient parasitic workers have been reared, they conduct well-organized raids of nearby host nests to capture their brood (D’Ettorre and Heinze, 2001). Some brood is eaten, but most workers eclose in the parasite’s nest and contribute to the workforce of the colony. By contrast, most inquiline species are tolerant of the host queen, allowing her to continuously produce host workers, whereas the inquiline queens focus their reproductive effort on sexual offspring (Hölldobler and Wilson, 1990; Bourke and Franks, 1991). Inquilines obligately depend on their hosts and most inquiline species lost their worker caste entirely (Kutter, 1968; Wilson, 1971, 1984; Hölldobler and Wilson, 1990).

The evolutionary origins of social parasitism have been debated since Darwin’s “Origin of Species” (Darwin, 1859). Entomologists have long noticed that ant social parasites and their hosts are close relatives (Wasmann, 1891, 1902; Emery, 1909; Wheeler, 1901a,b, 1904), an observation subsequently referred to as “Emery’s rule” (Le Masne, 1956). Strictly interpreted, Emery’s rule postulates a sister group relationship between host and parasite, whereas a less restrictive or “loose” interpretation signifies for example a congeneric, but not necessarily a sister taxon relationship (Ward, 1989, 1996; Bourke and Franks, 1991; Rabeling et al., 2014). Consequently, two competing hypotheses were developed for explaining the speciation mechanisms of social parasites: (i) the interspecific hypothesis proposes that host and social parasite evolved reproductive isolation in allopatry, whereas (ii) the intraspecific hypothesis postulates that the social parasite evolved directly from its host in sympatry (Wheeler, 1919; Buschinger, 1970, 1986, 1990, 2009; Wilson, 1971; Ward, 1989, 1996; Bourke and Franks, 1991; Savolainen and Vepsäläinen, 2003; Rabeling et al., 2014; Degueldre et al., 2021). Empirical studies of temporary, dulotic, and host queen-intolerant workerless ant social parasites generally provide support for the interspecific hypothesis (Hasegawa et al., 2002; Kronauer et al., 2003; Janda et al., 2004; Beibl et al., 2005, 2007; Huang and Dornhaus, 2008; Maruyama et al., 2008; Heinze et al., 2015; Ward et al., 2015; Prebus, 2017; Sanllorente et al., 2018; Romiguier et al., 2018), whereas recent phylogenetic studies lend support to the intraspecific hypothesis for queen-tolerant inquilines (Savolainen and Vepsäläinen, 2003; Jansen et al., 2010; Rabeling et al., 2014; Leppänen et al., 2015; Nettel-Hernanz et al., 2015). In some cases, host shifts, secondary speciation events of hosts and/or parasites, and extinctions obscure the original evolutionary conditions under which social parasitism originated (Parker and Rissing, 2002; Krieger and Ross, 2005; Shoemaker et al., 2006).

To explore the origin and evolution of diverse socially parasitic life histories in eusocial insects, we reconstructed the evolutionary history of the Holarctic ant genus *Formica. Formica* ants are ideally suited for comparative studies of social parasitism because the genus has the highest number of social parasite species in any ant genus (87 out of 176; Table 1), and all socially parasitic life history traits known from eusocial insects evolved in *Formica* ants (Figure 1, Tables 1 and 2). In addition, colonies of *Formica* species vary significantly in colony founding behavior as well as in nest and colony structures, providing an opportunity to explore the interplay between colony organization and life history at the origin of social parasitism. Some *Formica* species use independent colony foundation, when new colonies are started by a single queen (i.e., haplometrosis) or a group of cooperating queens (i.e., pleometrosis). Queens of other species rely on dependent colony founding, cooperating with groups of conspecific workers to found a new colony or invading an existing colony as a temporary or inquiline social parasites (Table 2) (Creighton, 1950; Letendre and Huot, 1972; Hölldobler and Wilson, 1990; Savolainen and Deslippe, 1996, 2001; Mori et al., 2001; Purcell et al., 2015; Brelsford et al., 2020). Furthermore, *Formica* colonies can have a single or multiple functional queens (monogyny vs. polygyny) and comprise one (monodomous) or multiple (polydomous) to thousands interconnected physical nests covering a large area (supercolonial) (Helanterä et al., 2009).

**Table 1:** Diversity of social parasites in the genus *Formica* compared to all other ants. Socially parasitic life histories are significantly overrepresented in *Formica* ants, except for inquilinism. The total social parasite diversity in ants is higher than the sum of species in individual life history categories because the biology of numerous social parasites remains unknown. The data are derived from published sources (Hölldobler and Wilson, 1990; Buschinger, 2009; Bolton, 2020; Rabeling, 2020).

|  | Temporary social parasites | Dulotic social parasites | Workerless inquiline social parasites | Total social parasite diversity |
| --- | --- | --- | --- | --- |
| All ants (n=13,861) | ~200 (1.4%) | ~80 (0.6%) | ~100 (0.7%) | >400 (2.9%) |
| *Formica* (n=176) | 72 (40.9%) | 14 (8.0%) | 1—2 (0.6—1.1%) | 87 (49.4%) |

**Table 2:**
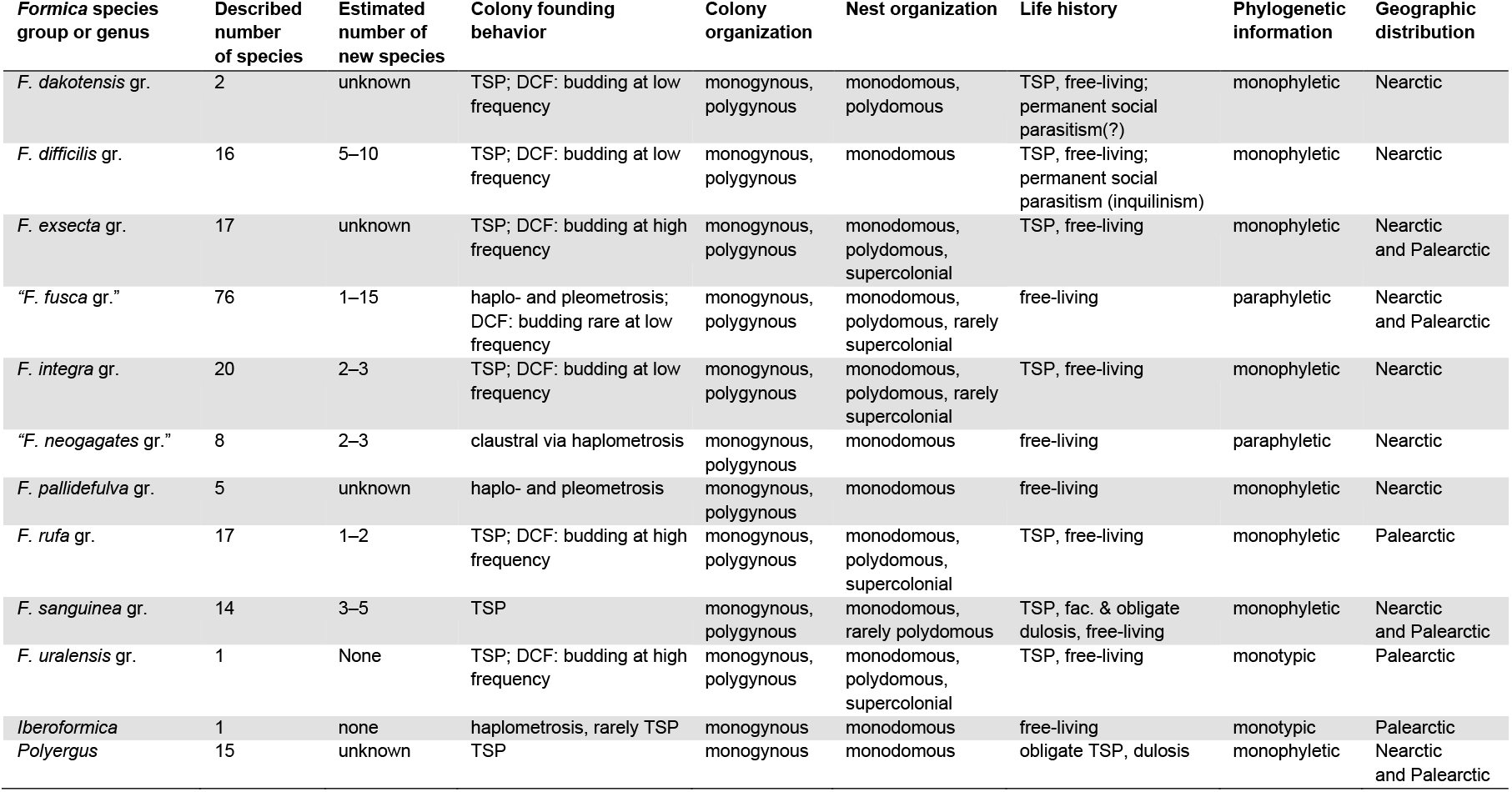
Diversity, taxonomy, life history, and evolutionary traits of *Formica* ants across currently recognized species groups, as well as of closely related formicine ants. The former *Formica rufa* group is divided into three clades, i.e., the *dakotensis, integra*, and *rufa* groups. The erstwhile *microgyna* group is properly referred to as the *difficilis* group based on name priority. Abbreviations: DCF: dependent colony founding; TSP: temporary social parasitism. Please refer to Table S2 for a detailed list of traits for individual species and references to original research. Total number of *Formica* species does not add to 176 because of one valid, poorly described species of uncertain group affinity.

**Figure 1:**
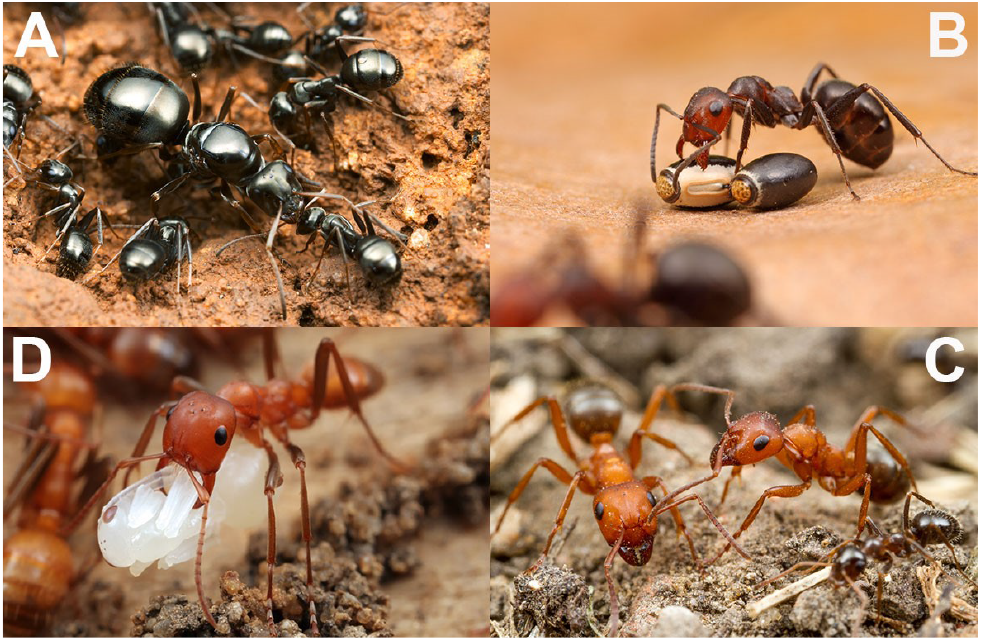
Diversity of life history traits in the formicine ants. In clock-wise direction: (A) members of the *Formica fusca* group practicing independent colony foundation; (B) *Formica obscuripes*, representing the *Formica integra* group (Nearctic members of the paraphyletic “*rufa*” group), which practices dependent and temporary social parasitic colony founding; (C) *Formica gynocrates*, representing the facultatively dulotic species of the *Formica sanguinea* group, with a worker of its *neogagates* group host species, *Formica vinculans*; (D) the highly modified worker of *Polyergus mexicanus*, representing the obligately dulotic formicine ants in the genera *Polyergus* and *Rossomyrmex*. All images courtesy of Alex Wild.

To infer the evolutionary origins of social parasitism and explore the behavioral transition from a cooperative social colony life to and exploitative socially parasitic life history, we reconstructed a global phylogeny for *Formica* ants and relevant outgroups from the formicine genera *Iberoformica, Polyergus, Proformica*, and *Rossomyrmex*, thus spanning the root node of the tribe Formicini (Blaimer et al., 2015). The comprehensive, time-calibrated phylogeny allows for (i) testing competing hypotheses regarding the origins and evolutionary transitions of social parasitism, (ii) reconstructing the evolutionary and biogeographic history of the group, and (iii) suggesting modifications to the internal classification of the genus.

## Results and discussion

### Formica originated in Eurasia during the Oligocene

To infer the life history evolution of the diverse, Holarctic genus *Formica*, we inferred a comprehensive phylogeny for 101 *Formica* species representing all ten currently recognized species groups (Table 2) across their wide geographic distribution in both the Old (19 spp.) and New World (82 spp.) and outgroups, using 2,242 Ultraconserved Element (UCE) loci per taxon. Our analyses recovered *Formica* as a strongly-supported clade with the monotypic genus *Iberoformica* as its sister lineage (Figure 2, Figure S1). *Formica* and *Iberoformica* split from their sister genus *Polyergus* around 33 Ma ago (95% Highest Posterior Density or HPD: 27–39 Ma) and *Formica* diverged from a common ancestor with its sister lineage *Iberoformica subrufa* approximately 30 Ma ago (95% HPD: 24–35 Ma). The crown group age of extant *Formica* ants is approximately 26 Ma (95% HPD: 21–31 Ma). Therefore, modern *Formica* ants evolved recently and likely originated in Eurasia during the Oligocene after the global cooling following the Terminal Eocene Event (Zachos et al., 2001). A similar evolutionary history was inferred for the species-rich Holarctic ant genus *Myrmica* (Jansen et al., 2010).

**Figure 2:**
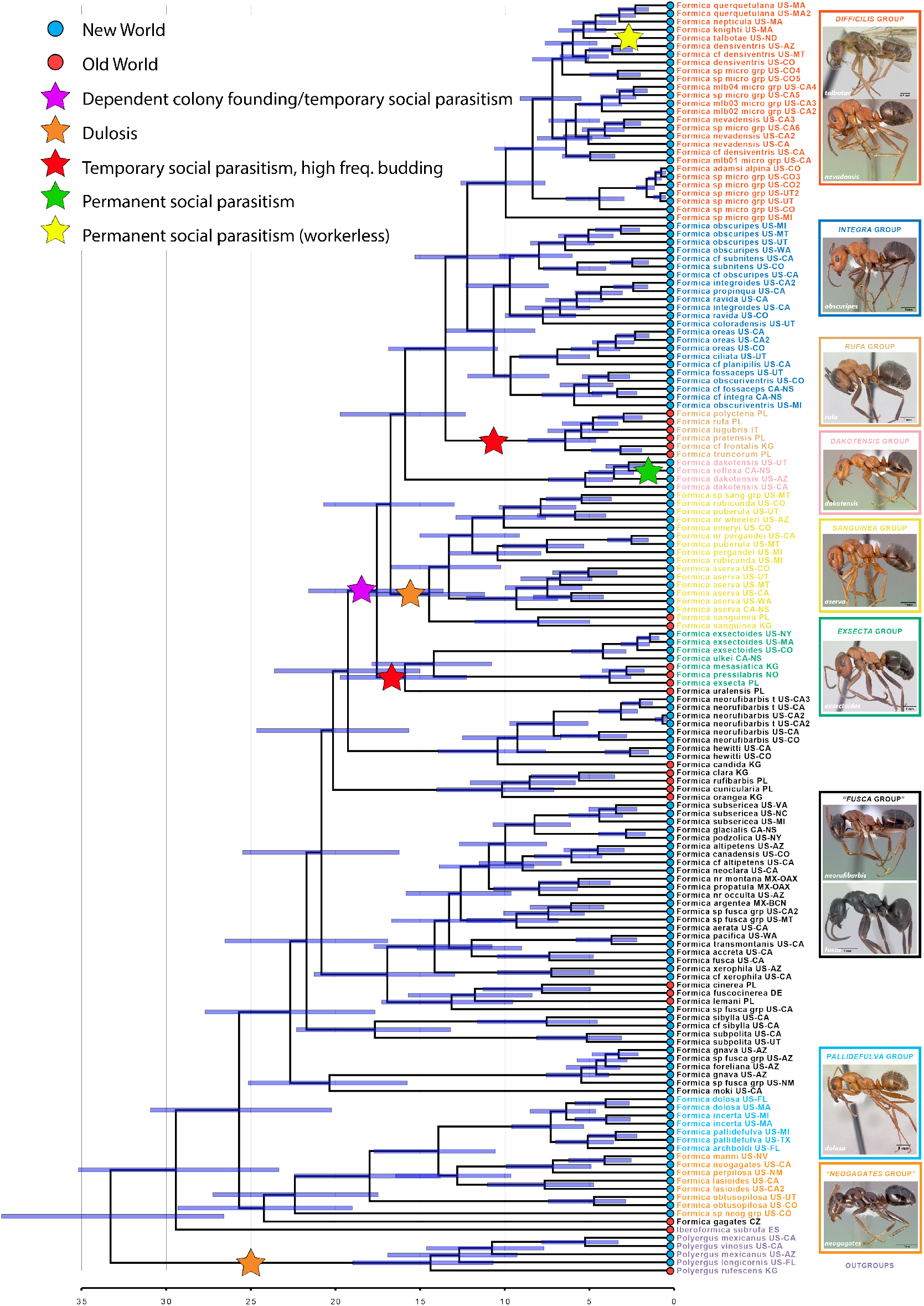
A time-calibrated molecular phylogeny of *Formica, Iberoformica* and *Polyergus*. Node bars indicate 95% highest posterior density. Scale in millions of years. Abbreviations after taxa names indicate sample origin. Country codes follow ISO 3166: USA (US), Canada (CA), and Mexico (MX); in addition to noting state/province. Note single origin of dependent colony foundation with inquilinism arising from within temporary parasites (yellow star). Taxon highlight colors signify species group membership; “*neogagates*” and “*fusca*” species groups are not monophyletic. Photographs by April Nobile and Erin Prado, courtesy of AntWeb.org

### Independent colony founding and social polymorphism are ancestral

To understand the evolution of social parasitism in *Formica* ants, it is necessary to recover the evolutionary origins of different life history strategies in the biologically diverse species groups. Character state reconstructions of life history traits including nest structure, social organization of colonies, and colony founding behaviors (Table 2, Figures S3–5) based on our phylogenomic tree (Figure 2) show that facultative polygyny and polydomy originated early during *Formica* evolution. Purely monogynous species groups are only found outside of *Formica* (Table 2). Within *Formica*, the deepest node marks the divergence between the independent colony founding species in the “*neogagates*” and *pallidefulva* groups plus *F. gagates* on one side and the independently colony founding species in the paraphyletic grade of the *Formica* “*fusca* group” on the other side of the bifurcation (Figure 2). Accordingly, facultative polygyny, independent colony founding, and polydomy are ancestral traits that were likely present in the most recent common ancestor (MRCA) of all extant *Formica* species. The “*fusca* group” arises as a paraphyletic grade nested between these early diverging lineages and all other *Formica* species and consists of at least five monophyletic groups (Figure 2). Transitions from independent to dependent colony founding behavior via colony budding evolved repeatedly in the “*fusca* group”, and independent colony founding behavior was completely lost when the ancestor of the clade including the *exsecta, sanguinea, dakotensis, rufa, integra* and *difficilis* species groups diverged from an ancestral “*fusca* group” member. The nest building behavior transitioned repeatedly between constructing monodomous, polydomous, and even supercolonial nests in the “*fusca* group”, which has important implications for population structure and population density of those species.

### Single origin of dependent and socially parasitic colony founding

In *Formica*, all socially parasitic and dependent colony founding lineages (Figures 1, 3) evolved secondarily from a shared single common ancestor practicing independent colony foundation approximately 18 Ma (95% HPD: 14–21 Ma). This clade of socially parasitic colony founding species includes species of *exsecta, sanguinea, dakotensis, rufa, integra* and *difficilis* groups, as well as *Formica uralensis*, a species of uncertain taxonomic affiliation which is here inferred as the sister lineage to the *exsecta* group. Recently, Romiguier and colleagues (Romiguier et al., 2018) also recovered a single origin of dependent colony foundation among Palearctic *Formica* species including representatives of four of the ten species groups. Our global phylogenetic analysis confirms and significantly expands on this earlier conclusion that both budding and parasitic colony founding evolved once, and adds a temporal scale showing that this event occurred around 18 Ma ago. Furthermore, our analysis reveals that clades of socially parasitic species have secondarily transitioned to other parasitic life histories. Evolutionary reversals from social parasitism to independent colony founding were not recovered, suggesting that a transition to a socially parasitic lifestyle is irreversible. A similar pattern was found in the ant genus *Lasius*, where temporary parasitism evolved twice but reversals to independent colony founding are unknown (Janda et al., 2004).

**Figure 3:**
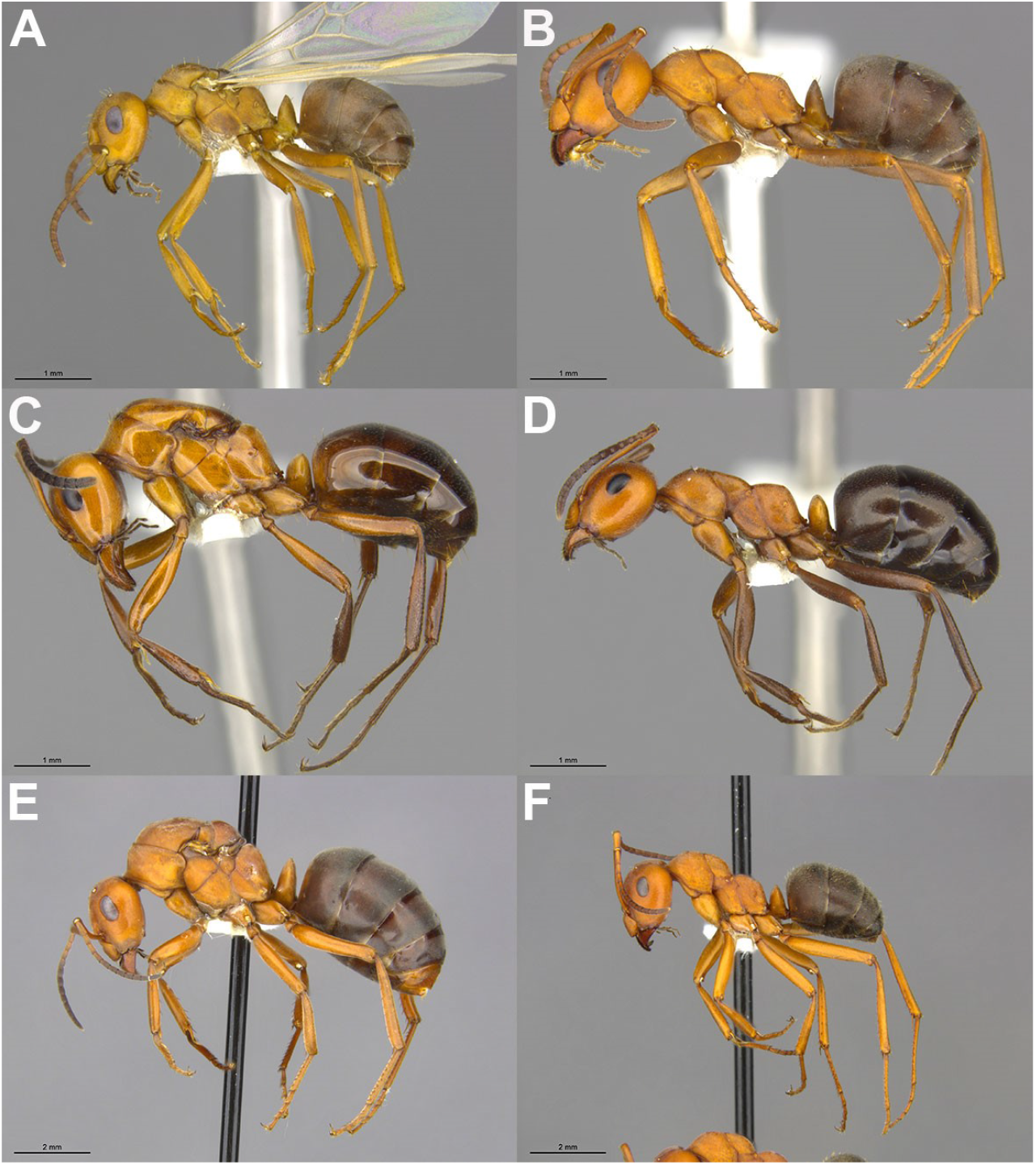
Convergent evolution of queen miniaturization in temporary social parasitic *Formica* ants. Queen miniaturization evolved in the *F. difficilis* (A, B) and *dakotensis* (C, D) species groups. In comparison, queens in the *integra* group (E, F) show a pronounced queen-worker dimorphism typical for *Formica* ants. Queen (A) and worker (B) of a hitherto undescribed *Formica difficilis* group species showing one of the most extreme cases of queen size reduction known in ants. Queen (C) and worker (D) of *Formica reflexa* representing a second, independent evolutionary origin of queen miniaturization, which is less extreme than in the *difficilis* group species. Queen (E) and worker (F) of *F. ravida* demonstrating a typically-sized queen with morphological modifications related to wing bearing and reproduction that are absent from the worker. Note that the *F. ravida* individuals are significantly larger than all other ants depicted in this figure. The scale bars represent 1 mm in figures A-D and 2 mm in figures E and F. Specimen identifiers are as follows A: MCZ 574034, B: MCZ 574022, C: MCZ 525288, D: MCZ 525283, E: MCZ 552096, F: MCZ 575163. Photographs by Patrick McCormack. Images copyright President and Fellows of Harvard University.

Among the socially parasitic colony founding species, two ecologically distinct life histories can be recognized. First, species in the *dakotensis, difficilis*, and *integra* groups are predominantly facultative temporary social parasites and practice colony budding at low frequency. Newly mated queens are unable to found new colonies independently via haplo- or pleometrosis, but instead, they seek adoption in heterospecific host colonies or re-adoption into conspecific colonies leading to secondary polygyny. In these species, colony budding seems to occur occasionally, which results in a characteristic population structure with smaller clusters of nests (usually less than 5), whereas large, unicolonial populations are absent. In contrast, species in the *exsecta* and *rufa* groups are facultative temporary social parasites practicing colony budding at high frequency, which can result in highly polydomous and/or unicolonial populations (Figure 2; Tables 2, S2).

The temporary social parasite species in the *Formica difficilis, integra, rufa*, and *dakotensis* groups shared a common ancestor approximately 16 Ma ago (95% HPD: 13–19 Ma), and they constitute the sister group to the dulotic *sanguinea* group (Figure 2). The obligate temporary social parasites in the *difficilis* group are also monophyletic and are sister to the *integra* group (Figure 2). All species in the *difficilis* group have miniature queens not larger than their largest workers (Figure 3a,b), which is likely associated with the socially parasitic life history. For the *difficilis* group, we infer a single origin of queen miniaturization and social parasitism that originated approximately 10 Ma ago (95% HPD: 8–12 Ma). It is important to note that most *difficilis* group species are rare and our knowledge about their biology is fragmentary at best. Therefore, the temporary social parasitic behavior remains to be observed for most species. However, the few existing direct observations on nest founding behavior, which include *F. difficilis* (Wheeler, 1904), *F. densiventris* (Wheeler, 1913), *F. impexa* (Wheeler, 1913), *F. adamsi alpina* (Cover, pers. obs.), and *Formica* new species (Cover, pers. obs.) confirm temporary social parasitism.

Interestingly, queen miniaturization evolved a second time, independently in the temporary social parasite species of the Nearctic *Formica dakotensis* clade (Figure 3), consisting of *F. dakotensis* and *F. reflexa* (Figure 2). *Formica dakotensis* is a facultative temporary social parasite (Creighton, 1950) and fully independent colonies are common. In contrast, *Formica reflexa* is rare and was only found in association with “*fusca* group” host workers (Buren, 1942; King and Sallee, 1951) (SP Cover, pers. obs.) suggesting a unique life history including parasitic colony founding and potentially a lifelong dependence on the host. Queen size reduction is frequently observed in inquiline social parasites (Bourke and Franks, 1991; Aron et al., 2004), but the independent origins of miniature queens in the *difficilis* and *dakotensis* clades imply that queen size reduction is adaptive for a temporary social parasitic life history syndrome in *Formica* ants.

### Evolution of dulosis

The dulotic species of the *Formica sanguinea* group are monophyletic (Figure 2), suggesting that dulotic behavior evolved once some time prior to its inferred crown group age of approximately 14 Ma (95% HPD: 11– 18 Ma). Thus, dulotic behavior and temporary social parasitism did not evolve simultaneously in *Formica*, but instead dulosis evolved secondarily from a temporary socially parasitic ancestor. The single origin of dulotic behavior in a diverse clade of temporary social parasite species supports the hypothesis that dulosis originates only under rare circumstances (D’Ettorre and Heinze, 2001; Buschinger, 2009; Rabeling, 2020). In fact, the evolutionary origins of dulotic behavior in ants have been debated since Darwin’s “Origin of Species” (Darwin, 1859) and three not mutually exclusive hypotheses have been proposed to explain the origins of this highly specialized behavior: (i) predation, (ii) brood transport, and (iii) territorial competition (Darwin, 1859; Buschinger, 1970, 1986, 2009; Wilson, 1975; Alloway, 1980; Stuart and Alloway, 1982, 1983; Pollock and Rissing, 1989; Hölldobler and Wilson, 1990; D’Ettorre and Heinze, 2001).

Our phylogenetic results and behavioral observations indicate that the predatory behavior of temporary social parasites could lead to the evolution of facultative dulosis in *Formica*. Brood stealing would be favored by natural selection if the aid of heterospecific workers increased the parasite’s fitness, though we lack experimental studies demonstrating fitness benefits of kidnapped host workers to any *sanguinea* group species at present. Additional biological factors that were associated with the evolutionary origins of dulosis, including polygyny, polydomy, brood transport, and territoriality (Santschi, 1906; Wheeler, 1910; Buschinger, 1970; D’Ettorre and Heinze, 2001), can also be inferred for the common ancestor of the dulotic species in the *Formica sanguinea* group.

It is important to note that dulosis evolved convergently and under different ecological conditions in distantly related, non-predatory ants, such as the omnivorous, scavenging species in the genera *Temnothorax* and *Tetramorium* (Beibl et al., 2005, 2007; Heinze et al., 2015; Prebus, 2017; Feldmeyer et al., 2017; Alleman et al., 2018). This pattern suggests that alternative factors, such as territoriality and brood transport, likely play an important role in the origin of dulotic behavior in non-predatory ants. Across the ant tree of life, dulosis originated at least nine times convergently in distantly related clades (D’Ettorre and Heinze, 2001; Trager, 2013; Sanllorente et al., 2018), including three origins in the Formicini (Blaimer et al., 2015), and six origins in the Crematogastrini (Beibl et al., 2005; Ward et al., 2015; Prebus, 2017).

### Evolution of inquiline social parasitism

The only confirmed workerless inquiline social parasite in the genus *Formica* is *F. talbotae* (Talbot, 1977; Wilson, 1977). *Formica talbotae* is phylogenetically nested within the *difficilis* clade (Figure 2), suggesting that inquilinism evolved once from a facultatively polygynous ancestor practicing temporary social parasitism. To our knowledge this is the first empirical evidence for an evolutionary transition from temporary to workerless inquiline social parasitism; a hypothesis earlier suggested by Wilson (Wilson, 1971). *Formica dirksi* has also been repeatedly suggested to be a workerless social parasite of *F. subaenescens* in Maine (Wing, 1949; Buschinger, 2009; Ellison et al., 2012), but there is no natural history data substantiating this claim.

*Formica talbotae* is a distant relative of its *integra* group host, *F. obscuripes*, with which it shared a common ancestor approximately 12 Ma ago (95% HPD: 10–15 Ma). The host-parasite relationship of *F. talbotae* and *F. obscuripes* is consistent with the “loose” interpretation of Emery’s rule, where hosts and parasites can be congeners but not sister lineages, suggesting that *F. talbotae* evolved via the interspecific, allopatric route of social parasite evolution. This result contrasts with previous studies inferring workerless inquiline social parasites as directly evolving from free-living, closely related ancestors via the intraspecific, sympatric route of social parasite speciation (Savolainen and Vepsäläinen, 2003; Jansen et al., 2010; Rabeling et al., 2014; Leppänen et al., 2015). However, and in contrast to many host queen tolerant inquiline social parasites, *F. talbotae* was found exclusively in queenless host colonies. It seems unlikely that the miniature *F. talbotae* queen(s) assassinate the many and much larger host queens. Instead, it appears more likely that *F. talbotae* specializes on declining host colonies that lost their reproductive queen(s) (Talbot, 1977; Wilson, 1977). Preferentially inhabiting queenless host colonies is a highly specialized and rare behavior that was only described for few social parasite species (Bruch, 1928; Rabeling and Bacci, 2010; Schär and Nash, 2014; **?**). It is important to note that the phylogenetic placement of *F. talbotae* was not unequivocal in our analysis, but it is important for understanding the evolution of workerless inquilinism in *Formica* ants. The museum specimens of *F. talbotae* available to us were collected in the 1950s, and from these specimens we recovered only fragments of 496 UCE loci (∼8% of nucleotides in the full data matrix). Coalescent-based species tree estimation is known to suffer from missing data (Sayyari et al., 2017), and to test for incongruencies, we performed coalescent-based species tree estimations (Edwards et al., 2007; Edwards, 2009; Mirarab et al., 2014; Zhang et al., 2018) using a reduced data matrix that included only the 67 loci for which at least 50% of the *F. talbotae* sequences were present. This analysis recovered *F. talbotae* as the sister lineage to the *difficilis* group (Figure S8), and statistical support was low across the species tree. The phylogenetic position of *F. talbotae* differs from the nested position obtained in concatenation (Figure 2, Figure S1), but the topology is more similar to the concatenated tree than the species tree analysis of the full dataset (Figure S6), suggesting that missing data does have a negative impact on the species tree analysis. A quartet sampling analysis (Pease et al., 2018) revealed topological conflict at the same nodes where the concatenated tree differed from the species tree, but strongly supported monophyly of the *difficilis* group including *F. talbotae* nested within (Figure S9).

In agreement with morphological evidence, which places *F. talbotae* within the *difficilis* group (Talbot, 1977; Wilson, 1977), we also consider *F. talbotae* a member of the *difficilis* group. We interpret the placement outside of the *difficilis* group by species tree methods as an artifact caused by missing data.

### Historical biogeography

To infer the biogeographic conditions under which social parasitism evolved in the Formicini, we conducted a historic biogeography analysis inferring repeated dispersal between the Old and New World in the genera *Formica, Iberoformica*, and *Polyergus* (Figure 4, Figure S2). According to our biogeographical stochastic mapping analysis (Dupin et al., 2017), at least eight and more likely nine or ten such dispersal events occurred in this genus group (Figure 4). All other genera classified in the Formicini are confined to the Old World. Therefore, the genus *Formica* almost certainly originated in the Old World and subsequently started dispersing into the New World and back, likely via Beringia, which connected Eurasia and the Nearctic throughout most of the Paleogene (McKenna, 1983). Recent trans-Beringian dispersal was also demonstrated for several Holarctic ant species, which include *Formica gagatoides* (Schär et al., 2018).

**Figure 4:**
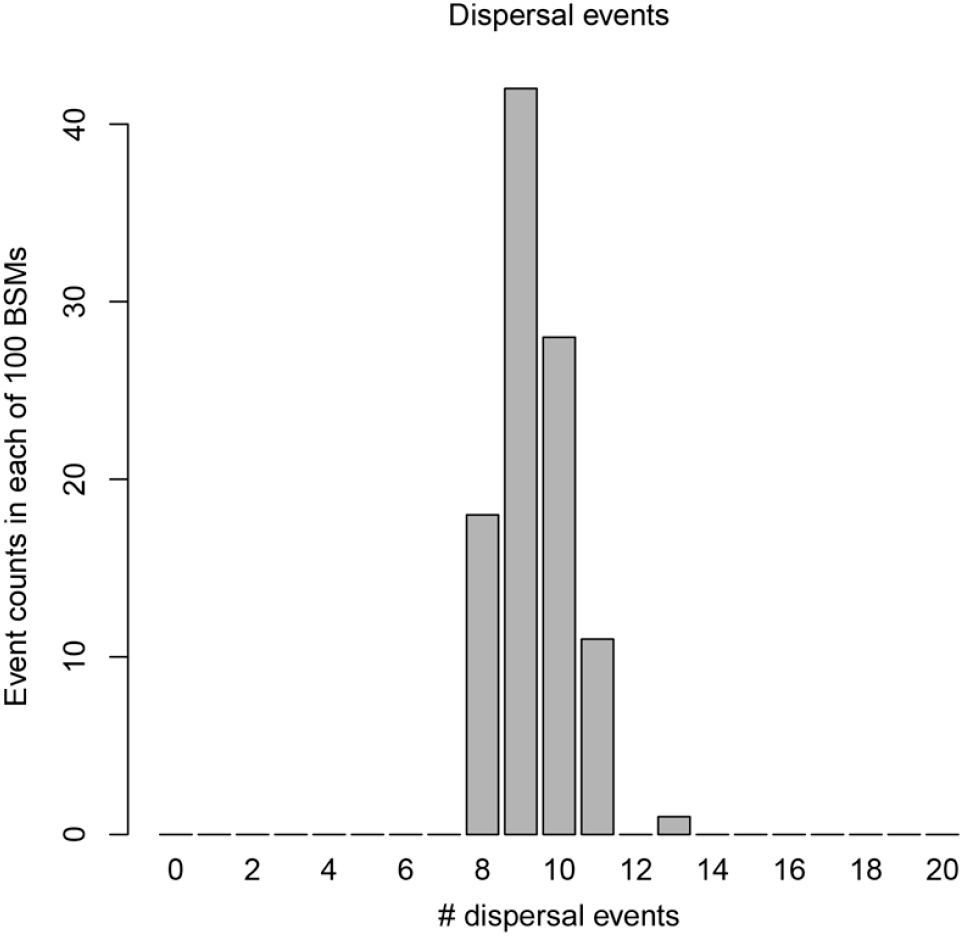
Inferred dispersal events between the Old and New World in *Formica, Iberoformica*, and *Polyergus* ants. Histogram of dispersal event counts from 100 biogeographic stochastic maps under the DEC+J model in BioGeoBEARS.

Nine fossil species of *Formica* are known from Eocene amber inclusions of Europe (47.8–33.9 Ma) (Aleksandrova and Zaporozhets, 2008a,b; Alekseev and Alekseev, 2016). Several fossils have been compared to extant species in the “*fusca*” and *rufa* groups (Wheeler, 1915), and close examination suggests that this resemblance may be superficial (Baroni Urbani and Graeser, 1987; Dlussky, 2008), which raises the question whether the Eocene amber fossils indeed represent crown group *Formica*? Doubts about the correct identification of putative *Formica* fossils in Baltic amber currently limit the utility of Eocene amber fossils for calibrating divergence analyses in the genus.

### Implications for Formica taxonomy and classification

The internal classification of species rich genera, such as *Formica*, is important because the affiliation with a species group (or subgenus) can provide first clues regarding the life history and general biology of an unknown species. Of the ten presently recognized *Formica* species groups (Table 2), the *pallidefulva* (= *Neoformica*), *exsecta, sanguinea* (= *Raptiformica*), and *difficilis* groups were recovered as monophyletic. In contrast, species traditionally classified in three species groups (or subgenera), the *fusca* (= *Serviformica*), *neogagates* (= *Proformica*), and *rufa* (= *Formica* s. str.) groups, form non-monophyletic assemblages. The Palearctic species *F. gagates*, traditionally placed in the *fusca* species group, was inferred as the sister species to the Nearctic clade comprising the *neogagates* and *pallidefulva* groups. The *neogagates* group itself forms a paraphyletic grade outside the *pallidefulva* group. The Holarctic *exsecta* group, plus the problematic *F. uralensis*, placed in a monotypic group, are sister to all other dependent colony founding/temporary social parasite groups, instead of *F. uralensis* being part of either the *fusca* or *rufa* groups, as previously suggested (Dlussky, 1967; Goropashnaya et al., 2012). The rest of species traditionally classified in *fusca* group form a grade consisting of five clades progressively more closely related to the dependently colony-founding clade. More thorough taxon sampling and a careful morphological study are necessary for a stable classification of those species. The Nearctic *F. obtusopilosa* is closely related to *F. neogagates* and not a member of the *sanguinea* group (Wilson and Brown, 1955; Buren, 1968), as treated by some authors (Gregg, 1963; Snelling and Buren, 1985). This classification is consistent with the biology of *F. obtusopilosa*, which is not dulotic and its queens found colonies independently, like other species in the *neogagates* group (Wilson and Brown, 1955; Gregg, 1963) (SP Cover, pers. obs.). Hence, the clypeal notch, long thought to be diagnostic of *sanguinea* group species, evolved convergently in *F. obtusopilosa*. The traditional *rufa* group is recovered as paraphyletic because the Old World *rufa* group species (now the true *rufa* species group) form a clade that is sister to the Nearctic *integra* and *difficilis* groups, the distinctive *F. dakotensis* and *F. reflexa* form a clade (here called the *dakotensis* group) sister to *rufa, integra*, and *difficilis* groups. Lastly, according to custom and for consistency we refer to the erstwhile *microgyna* group as the *difficilis* group, after the oldest constituent species name. The *difficilis* group species are monophyletic and sister to the *integra* group, not nested within it as previously suggested (Trager, 2016).

Considering that all the traditional *Formica* subgenera are nested within *Formica* and that three of the four subgenera are paraphyletic, we suggest discontinuing the use of the subgeneric names, and using species group names instead. While these results clarify the internal structure within the genus, much work remains to be done on the species level. The *fusca, integra, difficilis*, Nearctic *sanguinea* and *neogagates* groups all need taxonomic revisions.

### Conclusions

Our study provides a robust phylogenetic framework for studying the evolution of the diverse and ecologically important Holarctic ant genus *Formica* and allows for testing competing hypothesis regarding the origins and evolution of social parasitism in ants. We conclude that in the formicine genera *Formica, Polyergus*, and *Rossomyrmex*, social parasitism originated repeatedly and convergently. In the genus *Formica*, multiple transitions to increasingly more complex socially parasitic life histories evolved: (i) dependent colony foundation and facultative temporary social parasitism evolved once from an ancestor that established colonies via haplometrosis; (ii) dulosis, as practiced by members of the *sanguinea* group, evolved independently from dependent colony founding/facultatively temporary social parasitic ancestors; and (iii) the single known inquiline social parasite, *Formica talbotae*, evolved from a temporary social parasitic ancestor. Across species, *Formica* social parasites likely originated via the interspecific, allopatric speciation route of social parasite evolution, emphasizing that convergent evolutionary trajectories can lead to highly similar parasitic life history syndromes across eusocial insects.

The inferred sequence for the evolution of dulosis lends empirical support to Charles Darwin’s “predation hypothesis” for the origin of kidnapping behavior in ants. Furthermore, our results suggest that the ancestor of the dulotic *Formica* species likely possessed all the traits associated with the evolution of dulotic behavior, namely territorial and predatory behavior, brood transport behavior among spatially distinct nests of polygynous colonies, and the capacity for parasitic and dependent colony founding. The origin of dulosis was then followed by secondary diversification into 14 species which today form the monophyletic *sanguinea* group.

Our study inferred the workerless social parasite *Formica talbotae* as arising from a clade of temporary social parasites in the *difficilis* group, providing first empirical evidence for a transition from temporary to workerless social parasitism. The example of *Formica talbotae* underscores the importance for distinguishing between the different life history traits summarized under the umbrella term “inquiline social parasitism”. Taking the evolutionary origins into account is important because the majority of queen-tolerant workerless inquiline parasites likely speciated directly from free-living ancestors, whereas most queen-intolerant workerless social parasites apparently transitioned to the workerless state from a dulotic or temporary social parasitic ancestor. *Formica talbotae* can be regarded as a queen-intolerant obligate temporary social parasite that preferentially inhabits queenless host colonies and secondarily lost its worker caste. The distant relatedness to its host, *F. obscuripes*, and its distinct life history traits suggests that *F. talbotae* also evolved via the allopatric, interspecific route of social parasite evolution, contrasting with the sympatric origins of some queen-tolerant inquiline social parasites.

We show that *Formica* evolved during the early Oligocene, representing a relatively young ant genus that diversified rapidly into a diverse, ecologically dominant group. During its evolutionary history *Formica* ants dispersed several times between the Old and New World.

Our study outlines the life history changes associated with the transition from eusociality to social parasitism in ants. Given the high diversity of social parasite species in the genus *Formica*, and considering the high degree of morphological and behavioral specialization, socially parasitic *Formica* species appear to be an ideally-suited study organism for investigating caste determination and for exploring the genetic basis underlying behavioral and life history evolution.

## Methods

For detailed methods see SI Appendix. Reads generates for this study are available at the NCBI Sequence Read Archive (to be submitted upon acceptance for publication). Other files used in analyses are available on Zenodo (DOI: 10.5281/zenodo.4341310).

We newly sequenced 101 ingroup morphospecies from all nine species groups of *Formica* ants that were recognized prior to our study and 8 outgroup species. Collection data associated with sequenced samples can be found in Table S1 and detailed voucher information is on Zenodo.

To obtain the genetic data we extracted DNA, prepared genomic libraries and then enriched them using 9,446 custom-designed probes targeting 2,524 UCE loci in Hymenoptera (Branstetter et al., 2017). We submitted the enriched libraries to the University of Utah High Throughput Genomics Core Facility for sequencing on two Illumina HiSeq 125 Cycle Paired-End Sequencing v4 runs.

We processed the resulting reads using Phyluce bioinformatics pipeline (Faircloth, 2015). Following alignment and trimming we retained only individual locus alignments that had 110 or more taxa (70 % of total), resulting in 2,242 loci on average 667 nt long. The resulting concatenated matrix was 1,497,044 nt long and contained 17.58 % of missing data and gaps.

To infer the maximum likelihood phylogeny, we used ModelFinder (Kalyaanamoorthy et al., 2017) as implemented in IQ-TREE (Nguyen et al., 2015) to select the best model for each UCE locus under AICc. These models were then used for by-locus partitioned analysis of concatenated data matrix (Chernomor et al., 2016). To assess the robustness of this result to different analytics we performed an unpartitioned analysis and a quartet sampling analysis (Pease et al., 2018). In addition to concatenated analyses we performed coalescent-based species tree estimation using ASTRAL-III (Zhang et al., 2018).

For divergence time analyses we used a node dating approach, as implemented in MCMCTree, a part of the PAML package, v4.9e (Yang, 2007). We constrained our root node with soft bounds around a conservative maximum age estimate of 79 Ma, which corresponds to the lower bound of the 95 % highest posterior density interval for that split in a previous phylogenomic study (Blaimer et al., 2015).

For biogeographic inference we used BioGeoBEARS (Matzke, 2013). We discretized the distribution of *Formica* species into two regions, the New World and Old World. We used 100 replicates of biogeographical stochastic maps (Dupin et al., 2017) to estimate the number of times *Formica* dispersed between the Old and New World.

In order to investigate the evolution of nest structure, colony structure, and mode of colony foundation we used stochastic character mapping (Huelsenbeck et al., 2003) as implemented in the R package Phytools (Revell, 2011). We compared and selected best-fitting models of character evolution using GEIGER (Harmon et al., 2007) with time-calibrated tree pruned from distant outgroups and intraspecific samples as input. We based our character coding for each species (Table S2) on literature records and 80 years of cumulative field research by ourselves and colleagues.

## Supporting information

Suppmementary_Appendix

## Acknowledgments

This research was supported by the US National Science Foundation (NSF DEB-1456964, DEB-1654829, and NSF CAREER DEB-1943626). We gratefully acknowledge Philip Ward, James Trager, Matthew Prebus, Lech Borowiec, and André Francoeur for contributing important samples, as well as Jeffrey-Sosa Calvo, Benjamin Gerstner, and Cody Tipp for assisting with laboratory work and voucher specimen processing. Philip Ward and Jack Longino also contributed unpublished life history observations.

