## Supplementary material for "The evolution of social parasitism in *Formica* ants revealed by a global phylogeny": Suppmementary_Appendix

–

### Supplementary figures, tables, and references

Marek L. Borowiec
Stefan P. Cover
Christian Rabeling

#### Supplementary Methods

##### *Data availability*

Trimmed reads generated for this study are available at the NCBI Sequence Read Archive (to be submitted upon publication). Detailed voucher collection information, assembled sequences, analyzed matrices, configuration files and output of all analyses, and code used are available on Zenodo (DOI: 10.5281/zenodo.4341310).

##### *Taxon sampling*

For this study we gathered samples collected in the past ~60 years which were available as either ethanol-preserved or point-mounted specimens. Taxon sampling comprises 101 newly sequenced ingroup morphospecies from all seven species groups of *Formica* ants Creighton (1950) that were recognized prior to our study and 8 outgroup species. Our sampling was guided by previous taxonomic and phylogenetic work Creighton (1950); Francoeur (1973); Snelling and Buren (1985); Seifert (2000, 2002, 2004); Goropashnaya et al. (2004, 2012); Trager et al. (2007); Trager (2013); Seifert and Schultz (2009a,b); Muñoz-López et al. (2012); Antonov and Bukin (2016); Chen and Zhou (2017); Romiguier et al. (2018) and included representatives from both the New and the Old World. Collection data associated with sequenced samples can be found in Table S1.

##### *Molecular data collection and sequencing*

We performed non-destructive extraction and preserved same-specimen vouchers for each newly sequenced sample. We re-mounted all vouchers, assigned unique specimen identifiers (Table S1), and deposited them in the ASU Social Insect Biodiversity Repository (contact: Christian Rabeling,). Detailed voucher collection data can be accessed at 10.5281/zenodo.4341310. Briefly, we extracted DNA from all newly sequenced specimens using a DNeasy Blood and Tissue Kit (Qiagen, Valencia, CA, USA) and followed a library preparation protocol that follows was slightly modified from Blaimer et al. (2015). We used a KAPA Hyper Prep Library Kit (Kapa Biosystems, Inc., Wilmington, MA, USA) with magnetic bead cleanup Fisher et al. (2011) and a SPRI substitute Rohland and Reich (2012) as described in Faircloth et al. (2014). We used TruSeq adapters Faircloth and Glenn (2012) for ligation followed by PCR amplification of the library using a mix of HiFi HotStart polymerase reaction mix (Kapa Biosystems), Illumina TruSeq primers, and nuclease-free water.

We enriched each pool with 9,446 custom-designed probes (MYcroarray, Inc.) targeting 2,524 UCE loci in Hymenoptera Branstetter et al. (2017). We followed library enrichment procedures for the MYcroarray MYBaits kit Blumenstiel et al. (2010) except we used a 0.1× of the standard MYBaits concentration and added 0.7 µL of 500 µM custom blocking oligos designed against the custom sequence tags. We ran the hybridization reaction for 24 h at 65 °C, subsequently bound all pools to streptavidin beads (MyOne C1; Life Technologies) and washed bound libraries according to a standard target enrichment protocol Blumenstiel et al. (2010). We used the with-bead approach for PCR recovery of enriched libraries as described in Faircloth (2015).

We submitted pre-pooled libraries to the University of Utah High Throughput Genomics Core Facility for quality control on Agilent Bioanalyzer (Agilent Technologies, Santa Clara, CA, USA) and normalization. The pooled libraries were then sequenced using one full and one partial lane of a HiSeq 125 Cycle Paired-End Sequencing v4 run. Quality-trimmed sequence reads generated as part of this study are available from the NCBI Sequence Read Archive (to be submitted upon acceptance for publication).

*Processing of UCE data*

We performed read cleaning, assembly, and matching of assembled contigs to UCE probes using Phyluce bioinformatics pipeline Faircloth (2015). We trimmed the FASTQ data using Illumiprocessor, a wrapper around Trimmomatic Bolger et al. (2014), with default settings (LEADING:5, TRAILING:15, SLIDING-WINDOW:4:15, MINLEN:40). Assemblies were done using Trinity v20140717 Grabherr et al. (2011) with the phyluce_assembly_assemblo_trinity wrapper. We then assessed orthology by matching the assembled contigs to enrichment probe sequences with phyluce_assembly_match_contigs_to_probes (min_coverage=50, min_identity=80).

*Alignment*

We used phylogeny-aware UPP workflow Nguyen et al. (2015) to align all UCE sequences. We used AMAS Borowiec (2016) for alignment wrangling and obtaining summary statistics and AliView Larsson (2014) for visualization. Although alignment trimming has been criticized in the past Tan et al. (2015), we decided to trim the alignments because of substantial computational burden associated with analysis of untreated data with high proportion of gaps. We used trimAl Capella-Gutierrez et al. (2009) and its "gappyout" algorithm, which is a relatively relaxed algorithm for removal of gappy sites. Visual inspection of alignments revealed that occasionally sequences were misaligned towards flanks. To automatically identify and discard the misaligned sequences we wrote a custom R script (R Core Team) that leveraged packages ape Paradis et al. (2004); Paradis (2012), seqinr Charif and Lobry (2007), doParallel, and plyr. The script first generates a matrix of uncorrected p-distances from a UCE locus alignment and for each taxon it computes average p-distance to all other taxa. Then it creates a distribution of average per-locus p-distances for each taxon and detects outliers defined as sequences that lay above 3 SD from the mean of that distribution. Once identified, the script removes outliers using AMAS. This procedure resulted in removal of 0.97 % of all sequences. For downstream analyses we retained only alignments that had 110 or more taxa (70 % of total), resulting in 2,242 loci on average 667 nt long. The resulting concatenated matrix was 1,497,044 nt long and contained 17.58 % of missing data and gaps.

*Partitioning*

We used ModelFinder Kalyaanamoorthy et al. (2017) as implemented in IQ-TREE Nguyen et al. (2014). For each UCE we selected the best model under AICc. These models were then used for by-locus partitioned analysis of concatenated data matrix Chernomor et al. (2016). We have also employed the newly proposed strategy of partitioning UCE loci based on a sliding window approach that groups UCE sites with similar entropies Tagliacollo and Lanfear (2018). Unfortunately, many partitions identified using this approach were saturated and caused numerical instability in maximum likelihood analyses using IQ-TREE, resulting in unreasonably long tree lengths. We therefore proceeded with downstream analyses using per-locus partitioning.

*Phylogenetic and concordance analyses using maximum likelihood*

We used IQ-TREE Nguyen et al. (2014) for maximum likelihood inference of phylogeny on single-locus alignments and concatenated data matrix. To test the robustness of the partitioned concatenated analysis, we performed unpartitioned analysis under HKY+4G model, which was the most common model identified as best under AICc for single loci. To assess the sensitivity of our results using measures other than bootstrap support we performed a quartet sampling analysis with 500 maximum replicates (Pease et al., 2018). Results are summarized in Figure S7.

*Species tree analyses*

In addition to concatenated analyses we performed coalescent-based species tree estimation using ASTRAL-III (Zhang et al., 2018). Because summary coalescence methods such as ASTRAL have been shown to be negatively impacted by error in estimated gene trees Roch and Warnow (2015) we used the weighted statistical binning pipeline Mirarab et al. (2014); Bayzid et al. (2015). We collapsed all nodes with ultrafast bootstrap Minh et al. (2013) support below 95 for the binning pipeline, which resulted in identification of 1,733 supergenes containing from one to three UCE loci. We then used IQ-TREE to estimate supergene trees under a fully partitioned model (i.e. with branch lengths unlinked across partitions). Because of recent criticism of the statistical binning pipeline Adams and Castoe (2019) we also performed an analysis where raw trees from individual locus analyses were used. We mapped all terminals to putative species using morphology and the concatenated tree as guidance. Because some of the 101 species we recognized using morphology were non-monophyletic on the concatenated tree, we mapped the terminals onto 113 total monophyletic lineages representing putative species (Figures S4–S6). To test the effect of missing data Sayyari et al. (2017) on the position of *Formica talbotae* we performed additional analysis that used only the 67 loci which contained at least 50 % complete sequence for this taxon (Figure S6).

*Divergence time inference*

For divergence time analyses we used a node dating approach, as implemented in MCMCTree, a part of the PAML package, v4.9e Yang (2007). MCMCTree utilized rapid approximate likelihood computation dos Reis and Yang (2011), which makes it suitable for divergence dating of genome-scale data sets dos Reis et al. (2012). We constrained our root node with soft bounds around a conservative maximum age estimate of 79 Ma, which corresponds to the lower bound of the 95 % highest posterior density interval for that split in Blaimer et al. (2015). Although *Formica* has a rich fossil record, the affinity of these fossils is uncertain because, as this study shows, morphology has thus far been misleading about phylogeny. Because of this, we conservatively constrained the split of *Polyergus* and *Formica*+*Iberoformica* to be at least 34 Ma, or one of the younger estimates for the age of Baltic amber Aleksandrova and Zaporozhets (2008a,b). We ran each analysis unpartitioned, under the HKY+4G model for 20 million generations. We examined each run's statistics in Tracer.

*Biogeographic analyses*

For biogeographic inference we used BioGeoBEARS v1.1.2 Matzke (2013). We discretized the distribution of *Formica* species into two regions, the New World and Old World. Model selection implemented in BioGeoBEARS suggested DEC+J Matzke (2014) as the best-fitting model and we used it for all downstream analyses. We used 100 replicates of biogeographical stochastic maps Dupin et al. (2017) to estimate the number of times *Formica* dispersed between the Old and New World. We used our time-calibrated tree with taxon duplicates removed such that each species or putative species was represented by only one terminal. Results are summarized in Figure S2.

*Ancestral state reconstruction*

In order to investigate the evolution of natural history traits in *Formica* and *Polyergus* we used stochastic character mapping (Huelsenbeck et al., 2003). We used the same time-calibrated tree as for biogeographic analyses but with distant (*Rossomyrmex minuchae and Proformica mongolica*) outgroups and *Formica* taxa for which we had insufficient life history data (see Table S2) pruned.

For each species we collected literature and previously unpublished field research data on nest structure, colony structure, and colony foundation mode. We discretized nest structure into three categories: i) monodomous, ii) polydomous, and iii) supercolonial. For the analysis we assigned the highest nest structure complexity recorded, meaning that if a species is known to form monodomous, polydomous, or supercolonial nests, we characterized it as supercolonial. We discretized colony structure into i) monogynous or ii) polygynous. We assigned species to the polygynous category if they have been observed to be either monogynous or polygynous. We discretized colony founding mode into five categories: i) independent colony foundation via haplometrosis or pleometrosis, queen readoption occurs, budding absent or at low frequency, ii) facultative temporary social parasitism with budding at low frequency, iii) facultative temporary social parasitism with budding at high frequency, iv) obligate temporary social parasitism and dulosis without budding, v) permanent social parasitism. The coding of character states for each species and references to original research articles are available in Table S2.

We performed ancestral state reconstruction on each of the three characters (nest structure, colony structure, and colony founding). We first compared three commonly-used variants of the discrete character evolution M*k* model (Lewis, 2001): all rates equal (character state change rates are equal for all states), symmetric transition rates (character state change rates are different for each pair of states), and all rates different (character state change rates are different for each transition). We also fit the meristic model (assuming characters change in step-wise fashion). We fit these four models using ”fit” functions the R package GEIGER v2.0.6 (Harmon et al., 2007) to all three characters. We used Akaike Information Criterion corrected for sample size (AICc) weights as computed by the ”aic.w” function in Phytools v0.6-44 (Revell, 2011) to see which model fit best. This approach identified the all rates different model to be the best fit for nest and colony structure and the symmetric model was found best-fitting for colony founding.

Ancestral reconstruction results are found in Figures S3-S5.

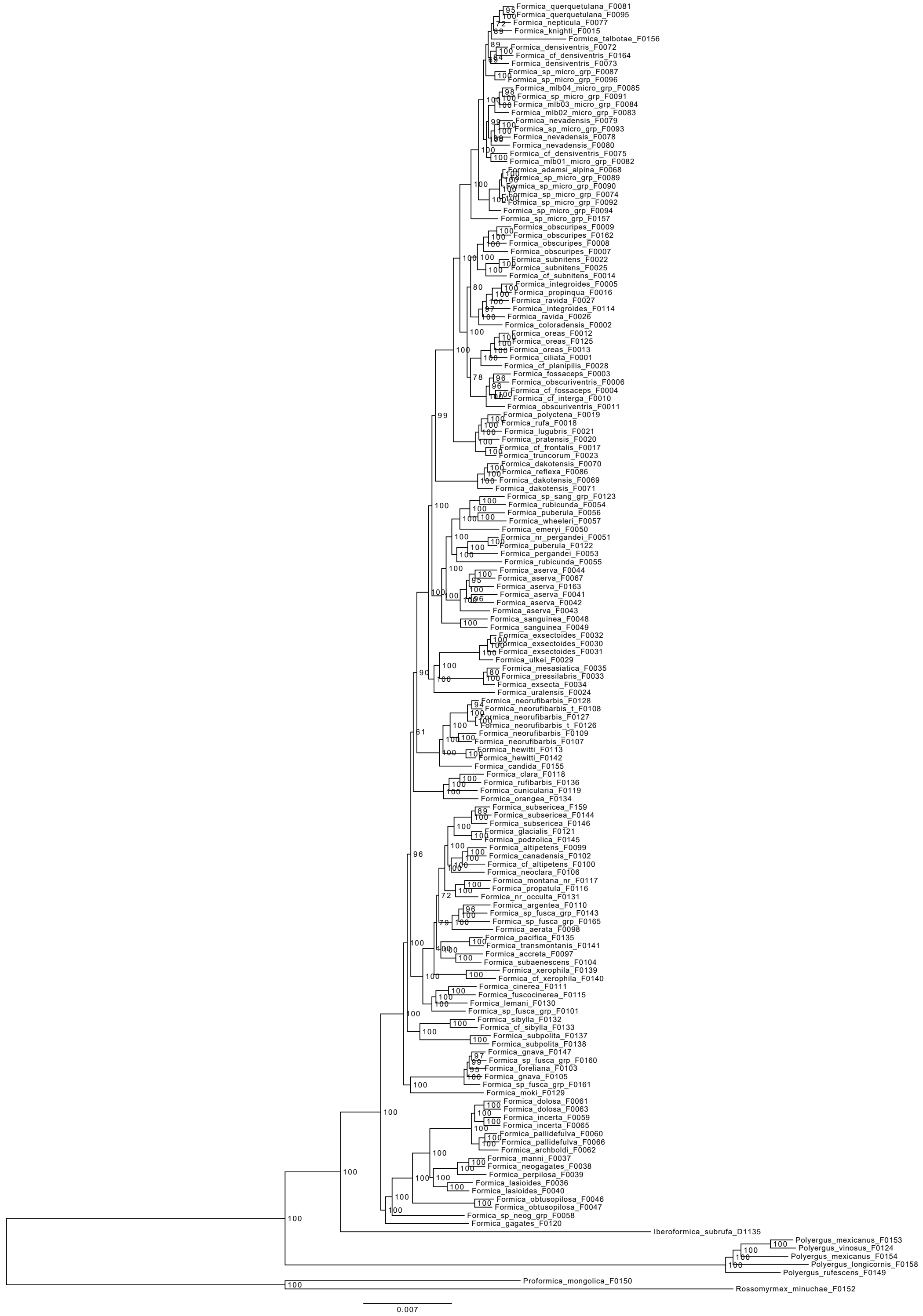


Supplementary Figure S1: Phylogram of *Formica* and outgroups from concatenated alignment inferred under maximum likelihood using IQ-TREE. Support in ultrafast bootstrap (values above 95 indicate strong support). Scale in substitutions per site.

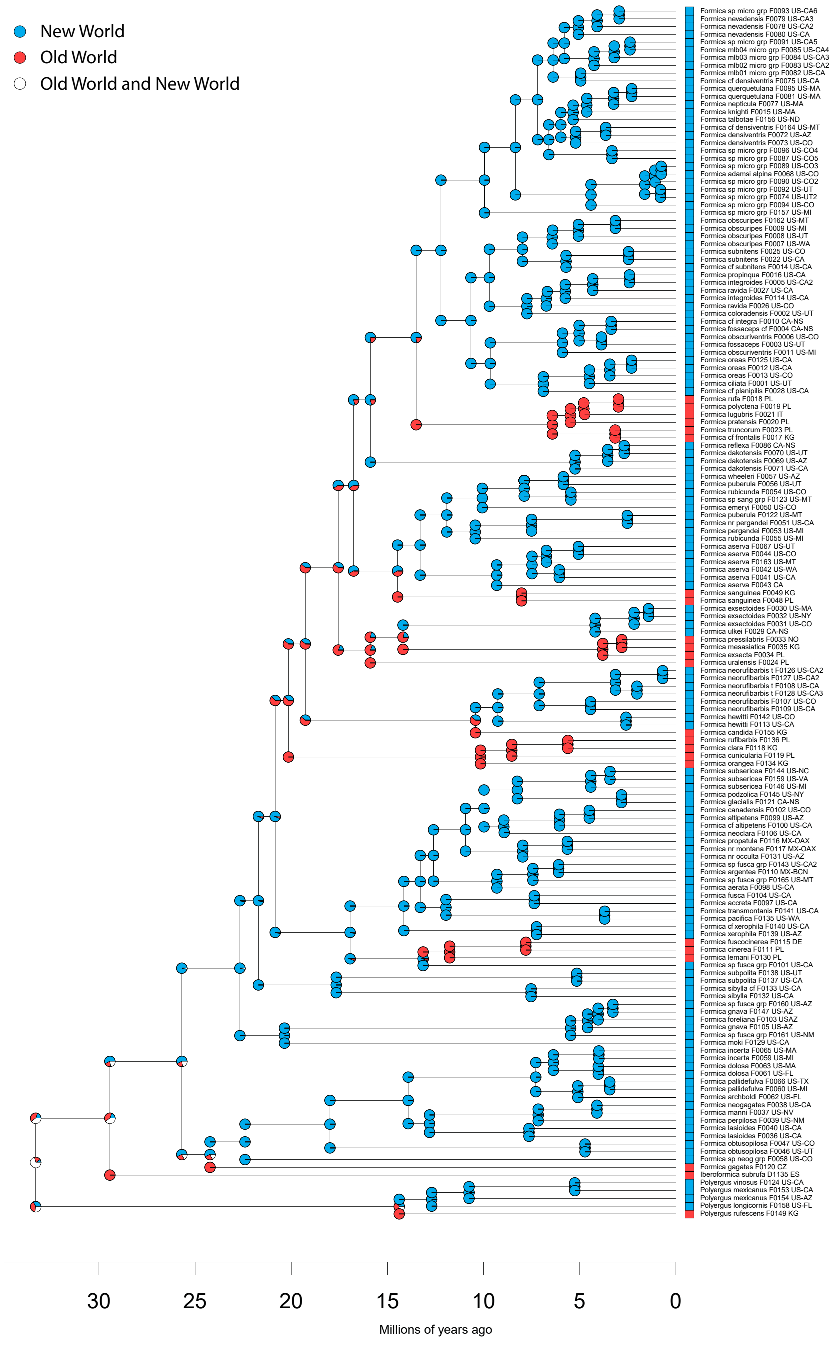


Supplementary Figure S2: Biogeographic history of *Formica*, *Iberoformica* and *Polyergus*. Pie charts depict relative likelihoods of range estimations from BioGeoBEARS under DEC+J. Pie charts at nodes correspond to ancestral state estimations and charts on the corners correspond to ranges immediately following speciation. Numbers preceded by the letter F indicate extraction code and correspond to identifiers in Table S1. Country and state/province codes as in main text Figure 1.

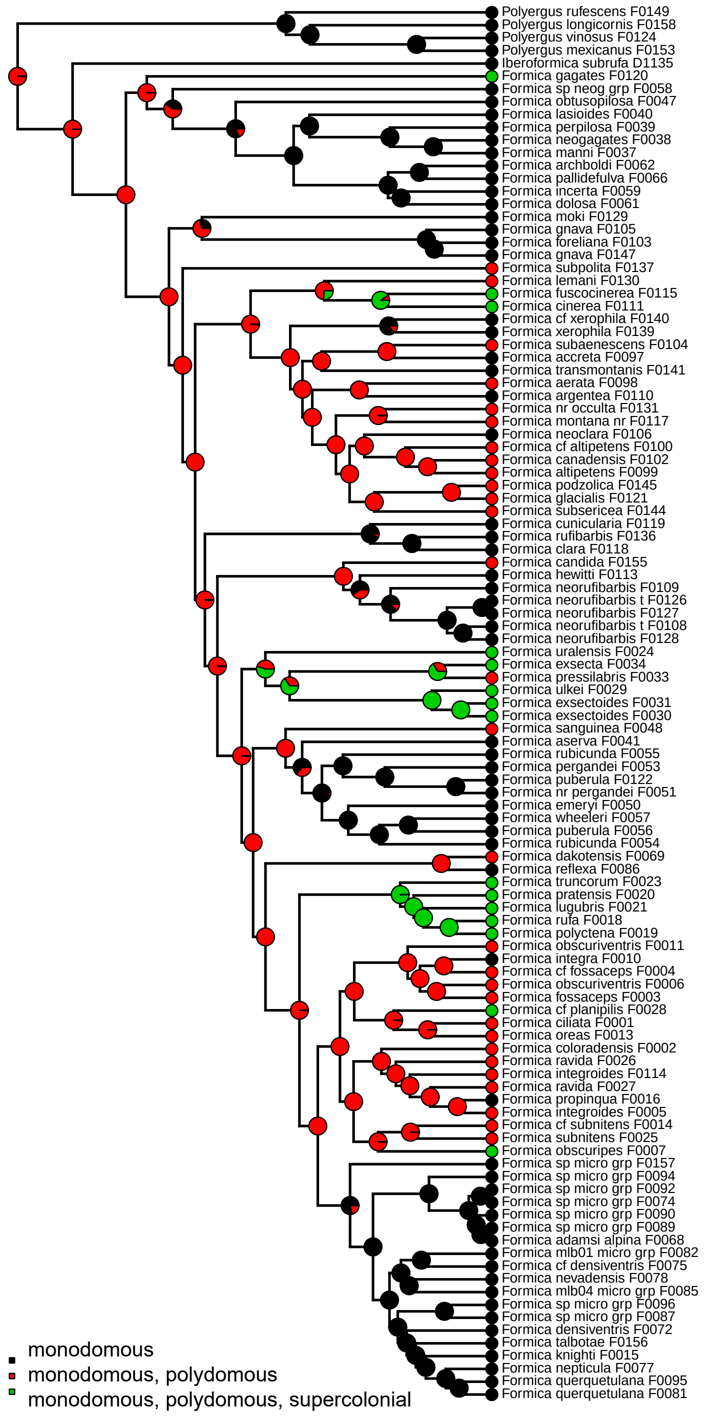


Supplementary Figure S3: Ancestral state reconstruction of nest structure using stochastic character mapping in Phytools of *Formica* nest structure under all rates different model. Pie charts at nodes correspond to ancestral state estimations and circles at tips correspond to states extant species. Numbers preceded by the letter F indicate extraction code and correspond to identifiers in Tables S1 and S2.

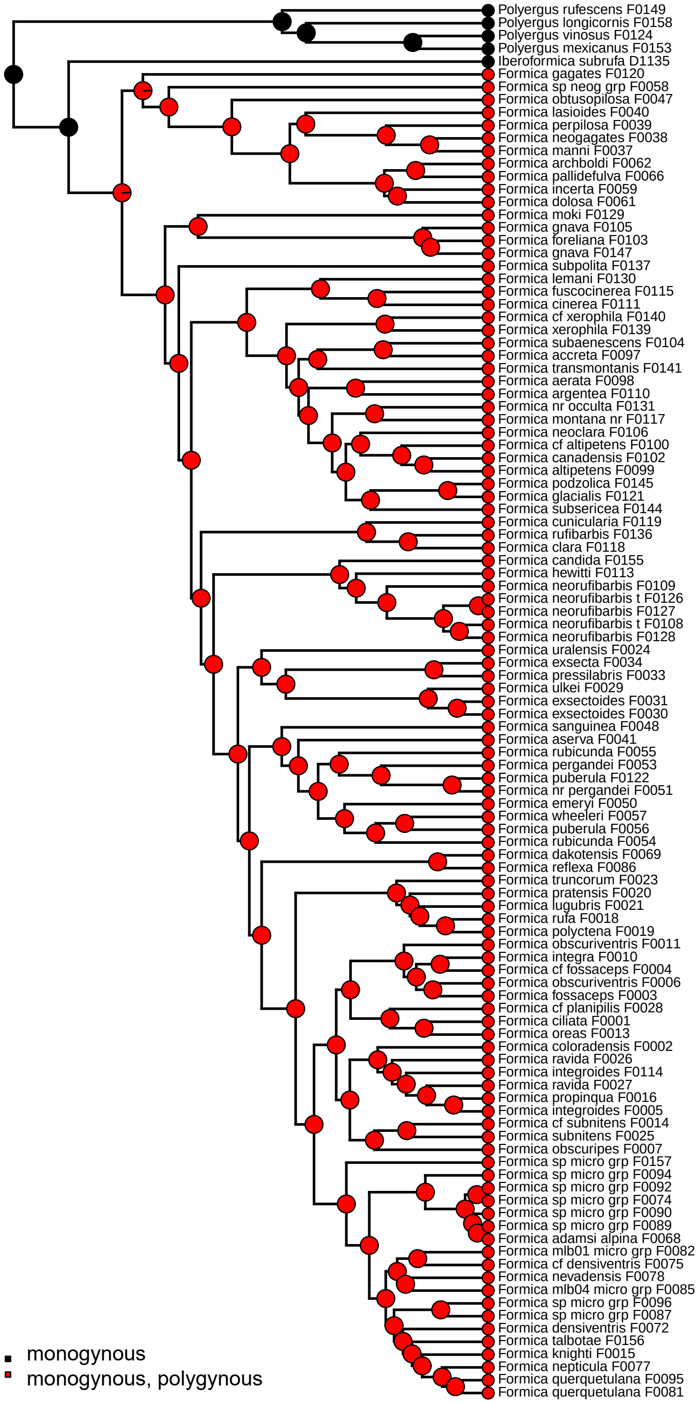


Supplementary Figure S4: Ancestral state reconstruction of colony structure using stochastic character mapping in Phytools of *Formica* colony structure under all rates different model. Pie charts at nodes correspond to ancestral state estimations and circles at tips correspond to states extant species. Numbers preceded by the letter F indicate extraction code and correspond to identifiers in Tables S1 and S2.

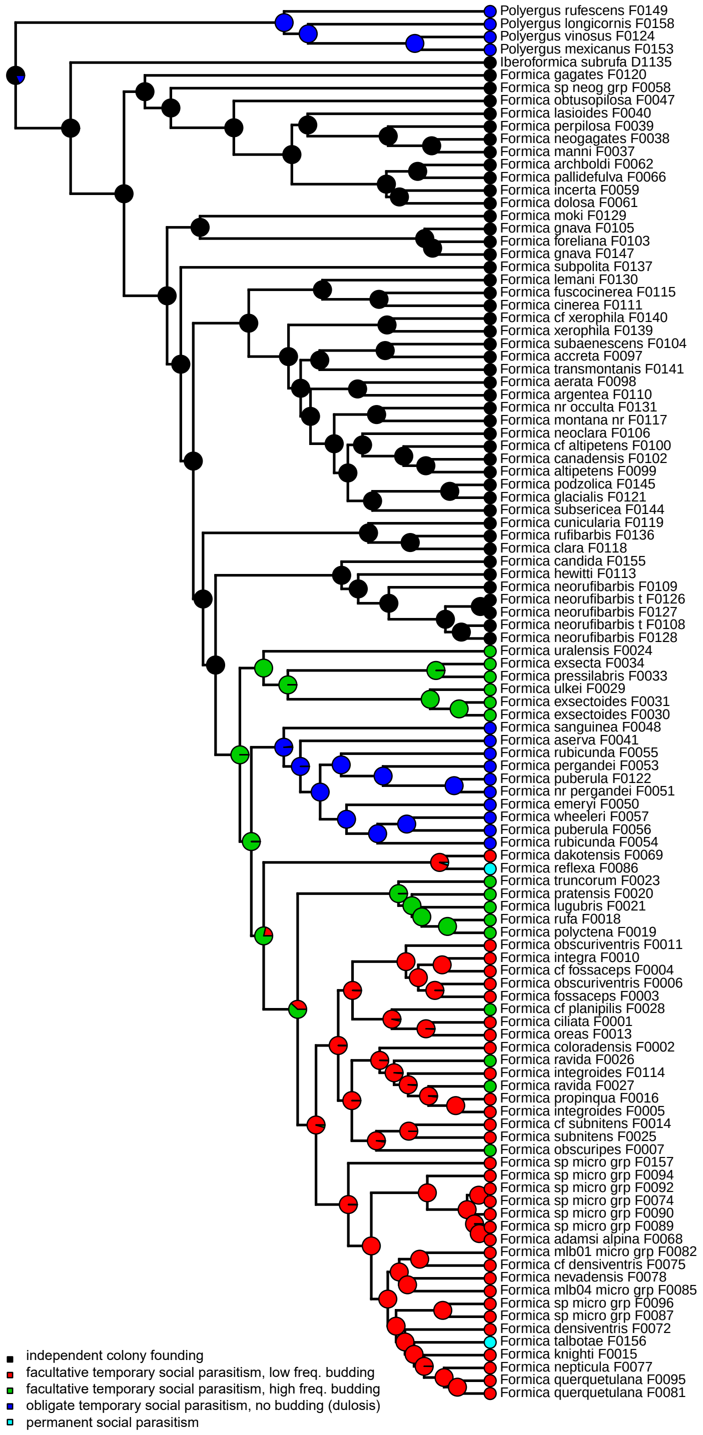


Supplementary Figure S5: Ancestral state reconstruction of colony founding mode using stochastic character mapping in Phytools of *Formica* colony founding under symmetrical rates model. Pie charts at the nodes correspond to ancestral state estimations and circles at tips correspond to states extant species. Numbers preceded by the letter F indicate extraction code and correspond to identifiers in Table S1 and S2.

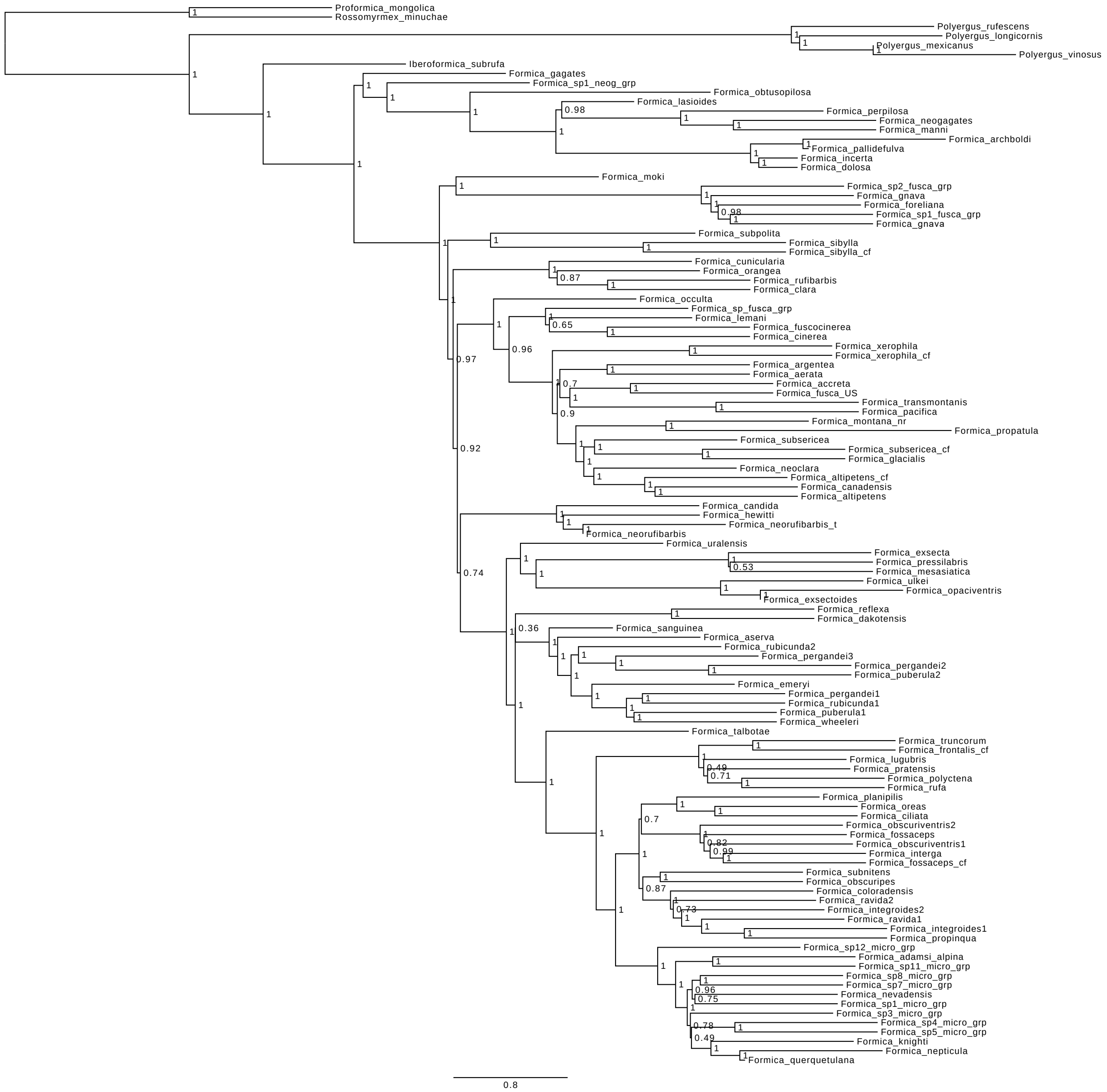


Supplementary Figure S6: Species tree inferred under coalescent using ASTRAL-III and 1,733 weighted supergene trees identified using statistical binning pipeline. Support is expressed in local posterior probability and branch lengths are in coalescent units.

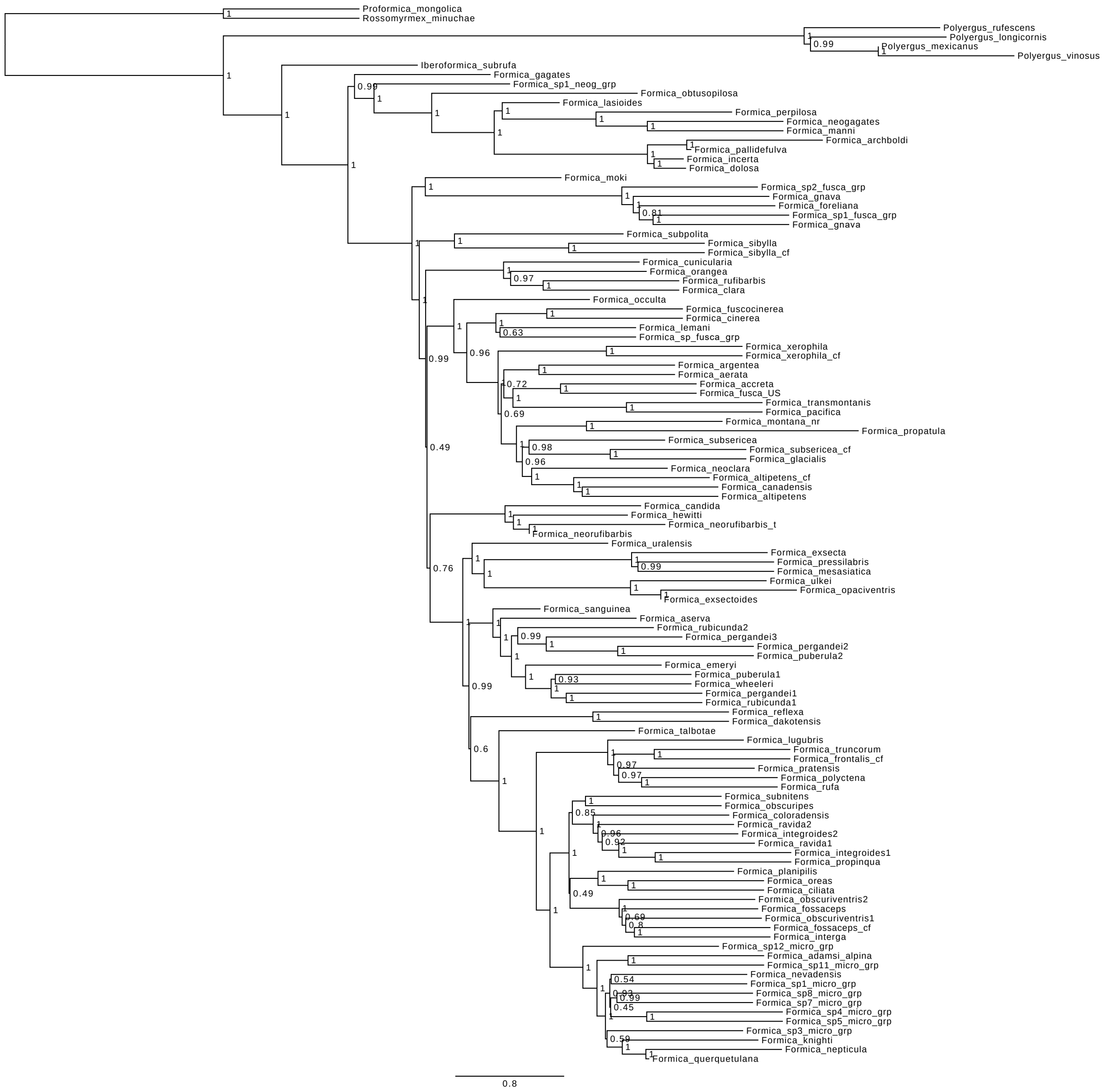


Supplementary Figure S7: Species tree inferred under coalescent using ASTRAL-III and all 2,242 individual UCE locus trees. Support is expressed in local posterior probability and branch lengths are in coalescent units.

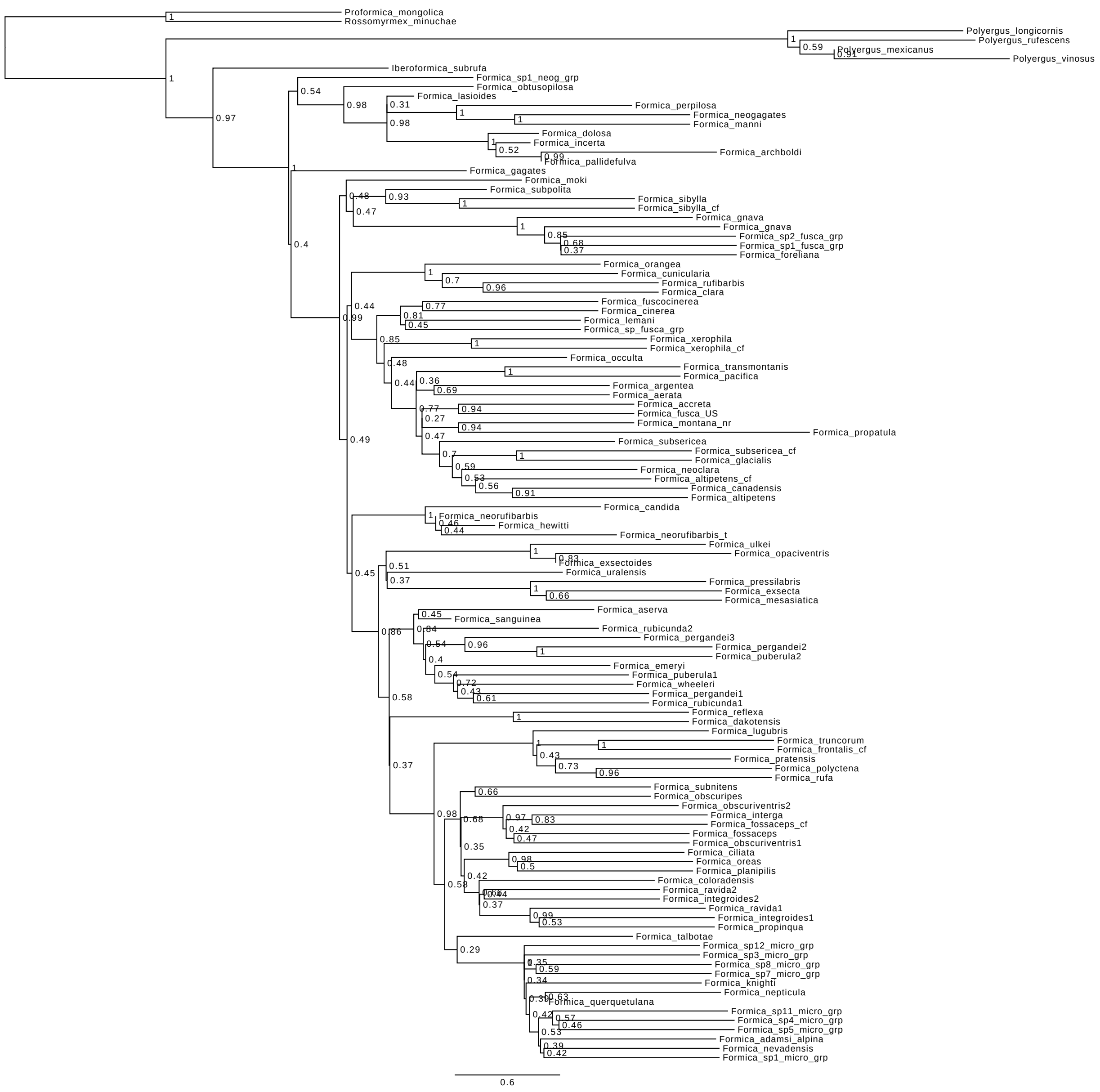

Supplementary Figure S8: Species tree inferred under coalescent using ASTRAL-III and 67 individual UCE locus trees for which at least 50 % of sequence length was available for *Formica talbotae*. Support is expressed in local posterior probability and branch lengths are in coalescent units.

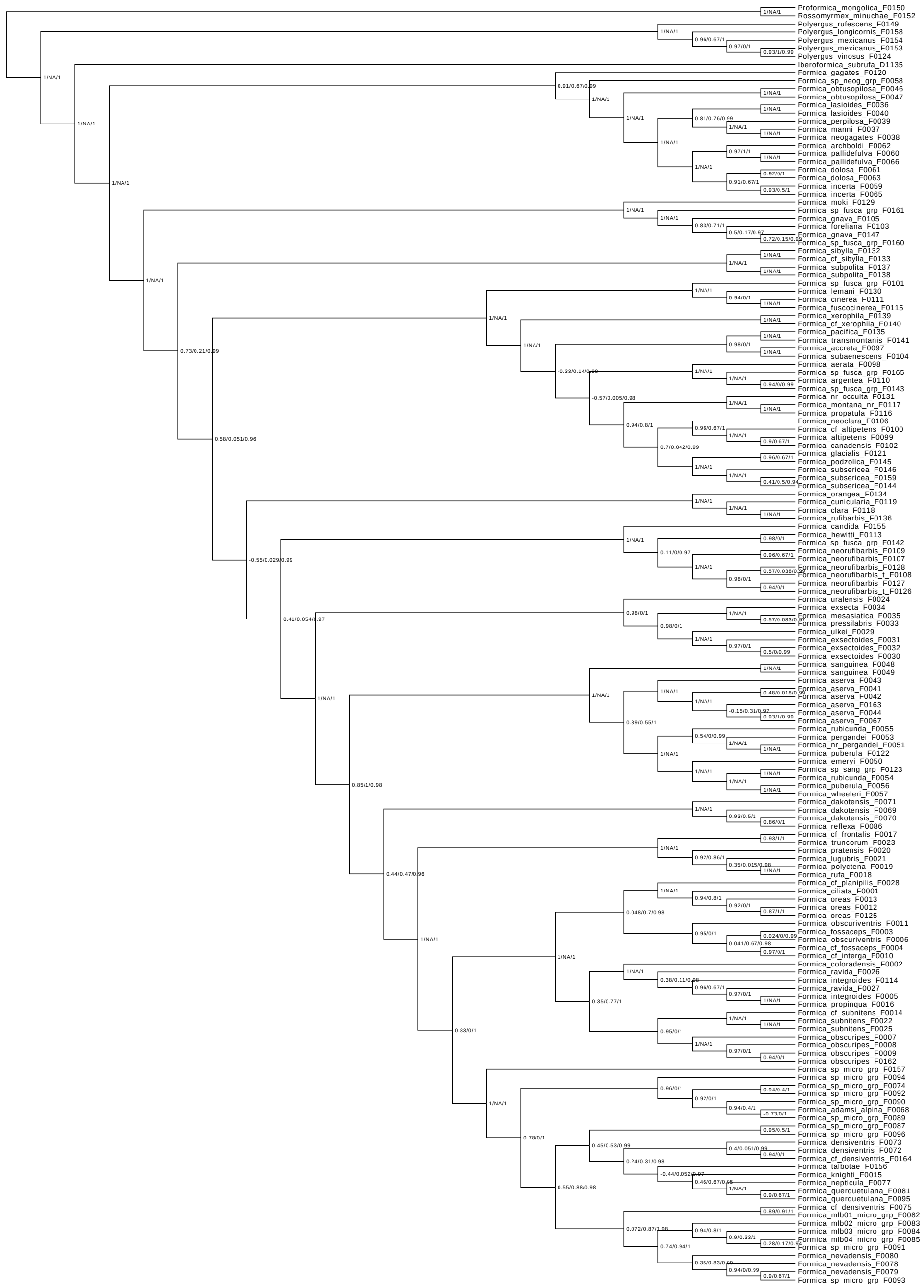


Supplementary Figure S9: Support from quartet sampling analyses. The values are: quartet concordance score (QC) / quartet differential score (QD) / quartet informativeness score (QI). Briefly, QC measures how often the concordant quartet topologies are inferred over discordant quartets in different replicates, with a value equal 1 when all replicates result in concordant topology, QD measures whether frequencies of two discordant topologies are equal (=1) or skewed (<1), and QI measures what proportion of replicates were informative. Contrast values for colony dependent clade or monophyly of *difficilis* group (1/NA/1), meaning that all replicates were informative and resulted in the same topology with support for placement of clade of Palearctic species related to *Formica rufibarbis* (-0.55/0.029/0.99) indicating that discordant topologies were inferred more often than those concordant with the input tree, that quartet topologies were highly skewed, but most replicates were informative. The former indicates maximal support, while the latter shows that the position of this clade within the ”fusca grade” is highly uncertain, perhaps due to incomplete lineage sorting.

Supplementary Table S1: Voucher specimens used in this study. Detailed collection data can be accessed at Zenodo (DOI: 10.5281/zenodo.4341310).

| Extraction ID | Taxon | Specimen ID | Country | Latitude | Longitude |
|---|---|---|---|---|---|
| D1135 | Iberoformica subrufa | ASU-SIBR2148 | ES | 37.185 | -3.485 |
| F0001 | Formica ciliata | ASU-SIBR2001 | US | 38.26117 | -112.51433 |
| F0002 | Formica coloradensis | ASU-SIBR2002 | US | 38.42233 | -109.18017 |
| F0003 | Formica fossaceps | ASU-SIBR2003 | US | 38.39467 | -109.165 |
| F0004 | Formica cf fossaceps | ASU-SIBR2004 | CA | 44.53050 | -64.31912 |
| F0005 | Formica integroides | ASU-SIBR2005 | US | 38.67525 | -119.99428 |
| F0006 | Formica obscuriventris | ASU-SIBR2006 | US | 39.12017 | -107.44767 |
| F0007 | Formica obscuripes | ASU-SIBR2007 | US | 48.1728 | -122.6714 |
| F0008 | Formica obscuripes | ASU-SIBR2008 | US | 38.23367 | -112.44917 |
| F0009 | Formica obscuripes | ASU-SIBR2009 | US | 42.44788 | -84.01688 |
| F0010 | Formica cf interga | ASU-SIBR2010 | CA | 44.68839 | -63.66808 |
| F0011 | Formica obscuriventris | ASU-SIBR2011 | US | 42.45103 | -84.01878 |
| F0012 | Formica oreas | ASU-SIBR2012 | US | 38.67577 | -119.99431 |
| F0013 | Formica oreas | ASU-SIBR2013 | US | 38.84233 | -106.53817 |
| F0014 | Formica cf subnitens | ASU-SIBR2014 | US | 38.67757 | -119.98343 |
| F0015 | Formica knighti | ASU-SIBR2015 | US | 41.8652 | -70.64777 |
| F0016 | Formica propinqua | ASU-SIBR2016 | US | 39.44008 | -120.32349 |
| F0017 | Formica cf frontalis | ASU-SIBR2017 | KG | 42.8 | 77.47 |
| F0018 | Formica rufa | ASU-SIBR2018 | PL | 51.155 | 16.985 |
| F0019 | Formica polyctena | ASU-SIBR2019 | PL | 51.39 | 17.5 |
| F0020 | Formica pratensis | ASU-SIBR2020 | PL | 51.3867 | 16.805 |
| F0021 | Formica lugubris | ASU-SIBR2021 | IT | 46.586 | 12.105 |
| F0022 | Formica subnitens | ASU-SIBR2022 | US | 39.82308 | -120.13748 |
| F0023 | Formica truncorum | ASU-SIBR2023 | PL | 50.33 | 16.55 |
| F0024 | Formica uralensis | ASU-SIBR2024 | PL | 51.38 | 23.55 |
| F0025 | Formica subnitens | ASU-SIBR2025 | US | 39.591 | -108.818 |
| F0026 | Formica ravida | ASU-SIBR2026 | US | 39.1695 | -107.94817 |
| F0027 | Formica ravida | ASU-SIBR2027 | US | 39.43221 | -120.24265 |
| F0028 | Formica cf planipilis | ASU-SIBR2028 | US | 40.72538 | -120.20708 |
| F0029 | Formica ulkei | ASU-SIBR2029 | CA | 44.49808 | -63.92692 |
| F0030 | Formica exsectoides | ASU-SIBR2030 | US | 41.87383 | -70.65183 |
| F0031 | Formica exsectoides | ASU-SIBR2031 | US | 38.73367 | -104.896 |
| F0032 | Formica exsectoides | ASU-SIBR2032 | US | 43.01548 | -77.57365 |
| F0033 | Formica pressilabris | ASU-SIBR2033 | SE | 68.3381 | 18.7639 |
| F0034 | Formica exsecta | ASU-SIBR2034 | PL | 50.577 | 22.984 |
| F0035 | Formica mesasiatica | ASU-SIBR2035 | KG | 42.33 | 78.24 |
| F0036 | Formica lasioides | ASU-SIBR2036 | US | 33.18522 | -116.28849 |
| F0037 | Formica manni | ASU-SIBR2037 | US | 41.63632 | -119.84183 |
| F0038 | Formica neogagates | ASU-SIBR2038 | US | 39.42998 | -120.24087 |
| F0039 | Formica perpilosa | ASU-SIBR2039 | US | 31.83945 | -109.03615 |
| F0040 | Formica lasioides | ASU-SIBR2040 | US | 39.43163 | -120.24059 |
| F0041 | Formica aserva | ASU-SIBR2041 | US | 37.68319 | -119.17078 |
| F0042 | Formica aserva | ASU-SIBR2042 | US | 48.66648 | -122.96887 |
| F0043 | Formica aserva | ASU-SIBR2043 | CA | 44.46380 | -63.58094 |
| F0044 | Formica aserva | ASU-SIBR2044 | US | 38.81400 | -106.28217 |
| F0046 | Formica obtusopilosa | ASU-SIBR2046 | US | 38.71217 | -111.94833 |
| F0047 | Formica obtusopilosa | ASU-SIBR2047 | US | 39.50633 | -108.76217 |
| F0048 | Formica sanguinea | ASU-SIBR2048 | PL | 50.448 | 19.995 |
| F0049 | Formica sanguinea | ASU-SIBR2049 | KG | 42.67 | 77.18 |
| F0050 | Formica emeryi | ASU-SIBR2050 | US | 39.03167 | -104.79633 |
| F0051 | Formica nr pergandei | ASU-SIBR2051 | US | 39.21529 | -121.04322 |
| F0053 | Formica pergandei | ASU-SIBR2053 | US | 42.45278 | -84.01939 |
| F0054 | Formica rubicunda | ASU-SIBR2054 | US | 39.03033 | -105.44217 |
| F0055 | Formica rubicunda | ASU-SIBR2055 | US | 42.45919 | -84.01429 |
| F0056 | Formica puberula | ASU-SIBR2056 | US | 37.88800 | -109.45433 |
| F0057 | Formica wheeleri | ASU-SIBR2057 | US | 31.91433 | -109.271 |
| F0058 | Formica sp neog grp | ASU-SIBR2058 | US | 37.84367 | -109.36967 |
| F0059 | Formica incerta | ASU-SIBR2059 | US | 42.45151 | -84.01901 |
| F0060 | Formica pallidefulva | ASU-SIBR2060 | US | 42.45948 | -84.0256 |
| F0061 | Formica dolosa | ASU-SIBR2061 | US | 30.35995 | -84.41848 |
| F0062 | Formica archboldi | ASU-SIBR2062 | US | 30.35995 | -84.41848 |
| F0063 | Formica dolosa | ASU-SIBR2063 | US | 41.81444 | -70.66315 |
| F0065 | Formica incerta | ASU-SIBR2065 | US | 41.8652 | -70.64777 |

Supplementary Table S1: Voucher specimens used in this study, continued.

| Extraction ID | Taxon | Specimen ID | Country | Latitude | Longitude |
|---|---|---|---|---|---|
| F0066 | Formica pallidefulva | ASU-SIBR2066 | US | 30.2275 | -98.1788 |
| F0067 | Formica aserva | ASU-SIBR2067 | US | 38.66283 | -111.94117 |
| F0068 | Formica adamsi alpina | ASU-SIBR2068 | US | 38.39317 | -107.19683 |
| F0069 | Formica dakotensis | ASU-SIBR2069 | US | 34.02867 | -109.1855 |
| F0070 | Formica dakotensis | ASU-SIBR2070 | US | 38.63500 | -111.94817 |
| F0071 | Formica dakotensis | ASU-SIBR2071 | US | 39.41344 | -120.32162 |
| F0072 | Formica densiventris | ASU-SIBR2072 | US | 32.65729 | -109.85841 |
| F0073 | Formica densiventris | ASU-SIBR2073 | US | 39.75867 | -108.78783 |
| F0074 | Formica sp micro grp | ASU-SIBR2074 | US | 38.64883 | -111.94983 |
| F0075 | Formica cf densiventris | ASU-SIBR2075 | US | 38.21541 | -119.74609 |
| F0077 | Formica nepticula | ASU-SIBR2077 | US | 41.87367 | -70.65233 |
| F0078 | Formica nevadensis | ASU-SIBR2078 | US | 39.35788 | -122.74716 |
| F0079 | Formica nevadensis | ASU-SIBR2079 | US | 41.56204 | -123.21177 |
| F0080 | Formica nevadensis | ASU-SIBR2080 | US | 39.41318 | -120.32038 |
| F0081 | Formica querquetulana | ASU-SIBR2081 | US | 41.87383 | -70.65183 |
| F0082 | Formica mlb01 micro grp | ASU-SIBR2082 | US | 37.21317 | -118.64671 |
| F0083 | Formica mlb02 micro grp | ASU-SIBR2083 | US | 40.94628 | -123.0427 |
| F0084 | Formica mlb03 micro grp | ASU-SIBR2084 | US | 39.29111 | -120.67918 |
| F0085 | Formica mlb04 micro grp | ASU-SIBR2085 | US | 39.40022 | -120.55818 |
| F0086 | Formica reflexa | ASU-SIBR2086 | CA | 44.53050 | -64.31912 |
| F0087 | Formica sp micro grp | ASU-SIBR2087 | US | 38.42033 | -107.62783 |
| F0089 | Formica sp micro grp | ASU-SIBR2089 | US | 38.18833 | -107.62067 |
| F0090 | Formica sp micro grp | ASU-SIBR2090 | US | 38.83800 | -106.56167 |
| F0091 | Formica sp micro grp | ASU-SIBR2091 | US | 38.83800 | -106.56167 |
| F0092 | Formica sp micro grp | ASU-SIBR2092 | US | 38.64983 | -111.95083 |
| F0093 | Formica sp micro grp | ASU-SIBR2093 | US | 41.56204 | -123.21177 |
| F0094 | Formica sp micro grp | ASU-SIBR2094 | US | 38.838 | -106.56167 |
| F0095 | Formica querquetulana | ASU-SIBR2095 | US | 41.87167 | -70.651 |
| F0096 | Formica sp micro grp | ASU-SIBR2096 | US | 39.04167 | -104.6615 |
| F0097 | Formica accreta | ASU-SIBR2097 | US | 39.30805 | -120.66831 |
| F0098 | Formica aerata | ASU-SIBR2098 | US | 37.80145 | -118.5299 |
| F0099 | Formica altipetens | ASU-SIBR2099 | US | 33.90683 | -109.1245 |
| F0100 | Formica cf altipetens | ASU-SIBR2100 | US | 38.6739 | -119.99411 |
| F0101 | Formica sp fusca grp | ASU-SIBR2101 | US | 37.38932 | -118.76612 |
| F0102 | Formica canadensis | ASU-SIBR2102 | US | 39.12017 | -107.44 |
| F0103 | Formica foreliana | ASU-SIBR2103 | US | 31.43100 | -111.16967 |
| F0104 | Formica subaenescens | ASU-SIBR2104 | US | 37.92051 | -122.57471 |
| F0105 | Formica gnava | ASU-SIBR2105 | US | 32.64983 | -109.81717 |
| F0106 | Formica neoclara | ASU-SIBR2106 | US | 39.50332 | -120.24157 |
| F0107 | Formica neorufibarbis | ASU-SIBR2107 | US | 38.71467 | -106.223 |
| F0108 | Formica neorufibarbis t | ASU-SIBR2108 | US | 36.67239 | -118.34463 |
| F0109 | Formica neorufibarbis | ASU-SIBR2109 | US | 39.42505 | -120.24285 |
| F0110 | Formica argentea | ASU-SIBR2110 | MX | 31.00261 | -115.5497 |
| F0111 | Formica cinerea | ASU-SIBR2111 | PL | 51.39 | 17.5 |
| F0113 | Formica hewitti | ASU-SIBR2113 | US | 38.67984 | -119.98373 |
| F0114 | Formica integroides | ASU-SIBR2114 | US | 40.11490 | -120.32086 |
| F0115 | Formica fuscocinerea | ASU-SIBR2115 | DE | 48.16452 | 11.50075 |
| F0116 | Formica propatula | ASU-SIBR2116 | MX | 17.1873 | -96.62085 |
| F0117 | Formica montana nr | ASU-SIBR2117 | MX | 17.61752 | -96.36799 |
| F0118 | Formica clara | ASU-SIBR2118 | KG | 42.66 | 77.47 |
| F0119 | Formica cunicularia | ASU-SIBR2119 | PL | 51.175 | 17.068 |
| F0120 | Formica gagates | ASU-SIBR2120 | CZ | 48.869 | 16.652 |
| F0121 | Formica glacialis | ASU-SIBR2121 | CA | 44.98735 | -64.06002 |
| F0122 | Formica puberula | ASU-SIBR2122 | US | 45.0446 | -110.68078 |
| F0123 | Formica sp sang grp | ASU-SIBR2123 | US | 45.13217 | -111.06283 |
| F0124 | Polyergus vinosus | ASU-SIBR2124 | US | 32.85323 | -116.43738 |
| F0125 | Formica oreas | ASU-SIBR2125 | US | 38.05273 | -119.32138 |
| F0126 | Formica neorufibarbis t | ASU-SIBR2126 | US | 38.06165 | -119.34966 |
| F0127 | Formica neorufibarbis | ASU-SIBR2127 | US | 38.06207 | -119.34888 |
| F0128 | Formica neorufibarbis | ASU-SIBR2128 | US | 38.04167 | -119.29781 |
| F0129 | Formica moki | ASU-SIBR2129 | US | 38.9 | -121.02 |
| F0130 | Formica lemani | ASU-SIBR2130 | PL | 50.33 | 16.55 |

Supplementary Table S1: Voucher specimens used in this study, continued.

| Extraction ID | Taxon | Specimen ID | Country | Latitude | Longitude |
|---|---|---|---|---|---|
| F0131 | Formica nr occulta | ASU-SIBR2131 | US | 31.91375 | -109.26757 |
| F0132 | Formica sibylla | ASU-SIBR2132 | US | 39.43221 | -120.24258 |
| F0133 | Formica cf sibylla | ASU-SIBR2133 | US | 39.431 | -120.331 |
| F0134 | Formica orangea | ASU-SIBR2134 | KG | 42.6 | 75.85 |
| F0135 | Formica pacifica | ASU-SIBR2135 | US | 48.1728 | -122.6714 |
| F0136 | Formica rufibarbis | ASU-SIBR2136 | PL | 52.76917 | 14.30778 |
| F0137 | Formica subpolita | ASU-SIBR2137 | US | 39.50444 | -120.23817 |
| F0138 | Formica subpolita | ASU-SIBR2138 | US | 38.72450 | -112.20717 |
| F0139 | Formica xerophila | ASU-SIBR2139 | US | 35.80350 | -112.11767 |
| F0140 | Formica cf xerophila | ASU-SIBR2140 | US | 38.85876 | -122.41834 |
| F0141 | Formica transmontanis | ASU-SIBR2141 | US | 39.188 | -123.757 |
| F0142 | Formica sp fusca grp | ASU-SIBR2142 | US | 38.83767 | -106.55933 |
| F0143 | Formica sp fusca grp | ASU-SIBR2143 | US | 39.21533 | -121.04019 |
| F0144 | Formica subsericea | ASU-SIBR2144 | US | 35.695 | -78.69688 |
| F0145 | Formica podzolica | ASU-SIBR2145 | US | 43.01616 | -77.57632 |
| F0146 | Formica subsericea | ASU-SIBR2146 | US | 42.45103 | -84.01885 |
| F0147 | Formica gnava | ASU-SIBR2147 | US | 31.93200 | -109.17783 |
| F0149 | Polyergus rufescens | ASU-SIBR2150 | KG | 42.67 | 77.18 |
| F0150 | Proformica mongolica | ASU-SIBR2151 | KG | 42.77 | 74.66 |
| F0152 | Rossomyrmex minuchae | ASU-SIBR2153 | ES | 36.89 | -2.78 |
| F0153 | Polyergus mexicanus | ASU-SIBR2154 | US | 39.41591 | -120.31704 |
| F0154 | Polyergus mexicanus | ASU-SIBR2155 | US | 31.90882 | -109.25211 |
| F0155 | Formica candida | ASU-SIBR2156 | KG | 42.08 | 76.73 |
| F0156 | Formica talbotae | ASU-SIBR2157 | US | 43.38 | -95.185 |
| F0157 | Formica sp micro grp | ASU-SIBR2158 | US | 42.45947 | -84.02559 |
| F0158 | Polyergus longicornis | ASU-SIBR2159 | US | 30.35995 | -84.41848 |
| F0159 | Formica subsericea | ASU-SIBR2160 | US | 38.9879 | -77.2495 |
| F0160 | Formica sp fusca grp | ASU-SIBR2161 | US | 31.8729 | -109.23504 |
| F0161 | Formica sp fusca grp | ASU-SIBR2162 | US | 32.59105 | -107.97398 |
| F0162 | Formica obscuripes | ASU-SIBR2163 | US | 48.03381 | -110.22798 |
| F0163 | Formica aserva | ASU-SIBR2164 | US | 46.7059 | -114.53617 |
| F0164 | Formica cf densiventris | ASU-SIBR2165 | US | 38.10233 | -111.33653 |
| F0165 | Formica sp fusca grp | ASU-SIBR2166 | US | 48.52302 | -113.3808 |

Supplementary Table S2: Life history of *Formica* and other taxa used in this study. Abbreviations: ASR: ancestral state reconstruction; ICF: independent colony founding; DCF: dependent colony founding; TSP: temporary social parasitism.

| Extraction ID | Taxon | Species group | Life history | Colony structure | Nest structure | Colony founding | ASR colony founding | References |
|---|---|---|---|---|---|---|---|---|
| F0097 | Formica accreta | "fusca" | free-living | monogynous, polygynous | monodomous | ICF | independent colony founding | Francoeur 1973; Frumhoff and Ward 1992; JT Longino and PS Ward unpubl. collection data |
| F0068 | Formica adamsi alpina | difficilis | free-living, TSP | monogynous, polygynous | monodomous | TSP, DCF: budding | facultative temporary parasitism, low frequency budding | SP Cover unpubl. collection data |
| F0098 | Formica aerata | "fusca" | free-living | monogynous, polygynous | monodomous, polydomous | ICF | independent colony founding | Francoeur 1973; PS Ward unpubl. collection data |
| F0099 | Formica altipetens | "fusca" | free-living | monogynous, polygynous | monodomous, polydomous | ICF, DCF: budding | independent colony founding | SP Cover unpubl. collection data |
| F0100 | Formica cf altipetens | "fusca" | free-living | monogynous, polygynous | monodomous, polydomous | ICF, DCF: budding | independent colony founding | SP Cover unpubl. collection data |
| F0062 | Formica archboldi | pallidevulva | free-living | monogynous, polygynous | monodomous | ICF: haplometrosis, pleometrosis | independent colony founding | Trager et al. 2007, SP Cover unpubl. collection data |
| F0110 | Formica argentea | "fusca" | free-living | monogynous, polygynous | monodomous | ICF: haplometrosis, pleometrosis | independent colony founding | Snyder 1992; SP Cover unpubl. collection data |
| F0041 | Formica aserva | sanguinea | free-living, TSP, fac. dulosis | monogynous, polygynous | monodomous | TSP | obligate temporary parasitism without budding (dulosis) | Savolainen and Deslippe 2001; SP Cover unpubl. collection data |
| F0042 | Formica aserva | sanguinea | free-living, TSP, fac. dulosis | monogynous, polygynous | monodomous | TSP | obligate temporary parasitism without budding (dulosis) | Savolainen and Deslippe 2001; SP Cover unpubl. collection data |
| F0043 | Formica aserva | sanguinea | free-living, TSP, fac. dulosis | monogynous, polygynous | monodomous | TSP | obligate temporary parasitism without budding (dulosis) | Savolainen and Deslippe 2001; SP Cover unpubl. collection data |
| F0044 | Formica aserva | sanguinea | free-living, TSP, fac. dulosis | monogynous, polygynous | monodomous | TSP | obligate temporary parasitism without budding (dulosis) | Savolainen and Deslippe 2001; SP Cover unpubl. collection data |
| F0067 | Formica aserva | sanguinea | free-living, TSP, fac. dulosis | monogynous, polygynous | monodomous | TSP | obligate temporary parasitism without budding (dulosis) | Savolainen and Deslippe 2001; SP Cover unpubl. collection data |
| F0163 | Formica aserva | sanguinea | free-living, TSP, fac. dulosis | monogynous, polygynous | monodomous | TSP | obligate temporary parasitism without budding (dulosis) | Savolainen and Deslippe 2001; SP Cover unpubl. collection data |
| F0102 | Formica canadensis | "fusca" | free-living | monogynous, polygynous | monodomous, polydomous | ICF, DFC: budding | independent colony founding | Francoeur 1973; SP Cover unpubl. collection data |
| F0155 | Formica candida | "fusca" | free-living | monogynous, polygynous | monodomous, polydomous | ICF | independent colony founding | Seifert 2004, Seifert 2018 |
| F0001 | Formica ciliata | integra | free-living, TSP | monogynous, polygynous | monodomous, polydomous | TSP, DCF: budding | facultative temporary parasitism, high frequency budding | Wheeler 1913; SP Cover unpubl. collection data |
| F0111 | Formica cinerea | "fusca" | free-living | monogynous, polygynous | monodomous, polydomous, supercolonial | ICF: haplometrosis, pleometrosis, DCF: budding | independent colony founding | Zhu et al. 2003; Lindström et al. 1996; Goropashnaya et al. 2001 |
| F0118 | Formica clara | "fusca" | free-living | monogynous, polygynous | monodomous | ICF | independent colony founding | Seifert 2007; Seifert 2018; Seifert and Schultz 2009 |
| F0002 | Formica coloradensis | integra | free-living, TSP | monogynous, polygynous | monodomous, polydomous | TSP, DCF: budding | facultative temporary parasitism, high frequency budding | Creighton 1940; SP Cover unpubl. collection data |
| F0119 | Formica cunicularia | "fusca" | free-living | monogynous, polygynous | monodomous | ICF: haplometrosis, pleometrosis | independent colony founding | Seifert and Schultz 2009; Seifert 2018 |
| F0069 | Formica dakotensis | dakotensis | free-living, TSP | monogynous, polygynous | monodomous, polydomous | TSP, DCF: budding | facultative temporary parasitism, high frequency budding | Wasmann 1902, Talbot 1972 |
| F0070 | Formica dakotensis | dakotensis | free-living, TSP | monogynous, polygynous | monodomous, polydomous | TSP, DCF: budding | facultative temporary parasitism, high frequency budding | Wasmann 1902, Talbot 1972 |
| F0071 | Formica dakotensis | dakotensis | free-living, TSP | monogynous, polygynous | monodomous, polydomous | TSP, DCF: budding | facultative temporary parasitism, high frequency budding | Wasmann 1902, Talbot 1972 |
| F0072 | Formica densiventris | difficilis | free-living, TSP | monogynous, polygynous | monodomous | TSP | facultative temporary parasitism, low frequency budding | Mackay and Mackay 2002; SP Cover and C Rabeling unpubl. collection data |
| F0073 | Formica densiventris | difficilis | free-living, TSP | monogynous, polygynous | monodomous | TSP | facultative temporary parasitism, low frequency budding | Mackay and Mackay 2002; SP Cover and C Rabeling unpubl. collection data |
| F0075 | Formica cf densiventris | difficilis | free-living, TSP | monogynous, polygynous | monodomous | TSP | facultative temporary parasitism, low frequency budding | SP Cover and C Rabeling unpubl. collection data |
| F0164 | Formica cf densiventris | difficilis | free-living, TSP | monogynous, polygynous | monodomous | TSP | facultative temporary parasitism, low frequency budding | SP Cover and C Rabeling unpubl. collection data |
| F0061 | Formica dolosa | pallidevulva | free-living | monogynous, polygynous | monodomous | ICF: haplometrosis, pleometrosis | independent colony founding | Trager et al. 2007, SP Cover unpubl. collection data |
| F0063 | Formica dolosa | pallidevulva | free-living | monogynous, polygynous | monodomous | ICF: haplometrosis, pleometrosis | independent colony founding | Trager et al. 2007, SP Cover unpubl. collection data |
| F0050 | Formica emeryi | sanguinea | TSP, dulosis | unknown | monodomous | TSP | obligate temporary parasitism without budding (dulosis) | Wheeler 1913; SP Cover unpubl. collection data |
| F0034 | Formica exsecta | exsecta | free-living, TSP | monogynous, polygynous | monodomous, polydomous, supercolonial | TSP, DCF: budding | facultative temporary parasitism, high frequency budding | Skibińska 1982; Pisarski 1982; Brown and Keller 2000; Haag-Liautard et al. 2008; Goryunov 2015 |
| F0030 | Formica exsectoides | exsecta | free-living, TSP | monogynous, polygynous | monodomous, polydomous, supercolonial | TSP, DCF: budding | facultative temporary parasitism, high frequency budding | Frumhoff and Ward 1992; SP Cover unpubl. collection data |
| F0031 | Formica exsectoides | exsecta | free-living, TSP | monogynous, polygynous | monodomous, polydomous, supercolonial | TSP, DCF: budding | facultative temporary parasitism, high frequency budding | Frumhoff and Ward 1992; SP Cover unpubl. collection data |
| F0032 | Formica exsectoides | exsecta | free-living, TSP | monogynous, polygynous | monodomous, polydomous, supercolonial | TSP, DCF: budding | facultative temporary parasitism, high frequency budding | Frumhoff and Ward 1992; SP Cover unpubl. collection data |
| F0103 | Formica foreliana | "fusca" | free-living | monogynous, polygynous | monodomous | ICF: haplometrosis | independent colony founding | Francoeur 1973; SP Cover unpubl. collection data |
| F0003 | Formica fossaceps | integra | free-living, TSP | monogynous, polygynous | monodomous, polydomous | TSP, DCF: budding | facultative temporary parasitism, high frequency budding | Buren 1942; SP Cover unpubl. collection data |
| F0004 | Formica cf fossaceps | integra | free-living, TSP | monogynous, polygynous | monodomous, polydomous | TSP, DCF: budding | facultative temporary parasitism, high frequency budding | SP Cover unpubl. collection data |
| F0017 | Formica cf frontalis | rufa | free-living, TSP | unknown | unknown | unknown | unknown | |
| F0115 | Formica fuscocinerea | "fusca" | free-living | monogynous, polygynous | monodomous, polydomous, supercolonial | ICF: haplometrosis, DCF: budding | independent colony founding | Seifert 2002; Seifert 2018 |
| F0120 | Formica gagates | gagates | free-living | monogynous, polygynous | monodomous, polydomous, supercolonial | ICF: haplometrosis, claustral | independent colony founding | Bernard 1967; Seifert 2018 |
| F0121 | Formica glacialis | "fusca" | free-living | monogynous, polygynous | monodomous, polydomous | ICF, DCF: budding | independent colony founding | Francoeur 1973; SP Cover unpubl. collection data |
| F0105 | Formica gnava | "fusca" | free-living | monogynous, polygynous | monodomous | ICF: haplometrosis | independent colony founding | Francoeur 1973; SP Cover unpubl. collection data |
| F0147 | Formica gnava | "fusca" | free-living | monogynous, polygynous | monodomous | ICF: haplometrosis | independent colony founding | Francoeur 1973; SP Cover unpubl. collection data |
| F0113 | Formica hewitti | "fusca" | free-living | monogynous, polygynous | monodomous | ICF | independent colony founding | Francoeur 1973; SP Cover unpubl. collection data |
| F0059 | Formica incerta | pallidevulva | free-living | monogynous, polygynous | monodomous | ICF: haplometrosis, pleometrosis | independent colony founding | Trager et al. 2007, SP Cover unpubl. collection data |
| F0065 | Formica incerta | pallidevulva | free-living | monogynous, polygynous | monodomous | ICF: haplometrosis, pleometrosis | independent colony founding | Trager et al. 2007, SP Cover unpubl. collection data |
| F0010 | Formica cf integra | integra | free-living, TSP | monogynous, polygynous | monodomous | TSP | facultative temporary parasitism, low frequency budding | Creighton 1940; SP Cover unpubl. collection data |
| F0005 | Formica integroides | integra | free-living, TSP | monogynous, polygynous | monodomous, polydomous | TSP | facultative temporary parasitism, low frequency budding | Creighton 1940; SP Cover unpubl. collection data |
| F0114 | Formica integroides | integra | free-living, TSP | monogynous, polygynous | monodomous, polydomous | TSP | facultative temporary parasitism, low frequency budding | Creighton 1940; SP Cover unpubl. collection data |
| F0015 | Formica knighti | difficilis | free-living, TSP | monogynous, polygynous | monodomous | TSP | facultative temporary parasitism, low frequency budding | Buren 1944; SP Cover and C Rabeling unpubl. collection data |
| F0036 | Formica lasioides | neogagates | free-living | monogynous, polygynous | monodomous | ICF: haplometrosis, claustral | independent colony founding | SP Cover unpubl. collection data |
| F0040 | Formica lasioides | neogagates | free-living | monogynous, polygynous | monodomous | ICF: haplometrosis, claustral | independent colony founding | SP Cover unpubl. collection data |
| F0130 | Formica lemani | "fusca" | free-living | monogynous, polygynous | monodomous, polydomous | ICF: haplometrosis, claustral | independent colony founding | Seppä et al. 2009; Gardner et al. 2007; Seifert 2018 |
| F0021 | Formica lugubris | rufa | free-living, TSP | monogynous, polygynous | monodomous, polydomous, supercolonial | TSP, DCF: budding | facultative temporary parasitism, high frequency budding | Pamilo et al. 1994; Maeder and Cherix 2001; Gyllenstrand and Seppä 2003 |
| F0037 | Formica manni | neogagates | free-living | monogynous | monodomous | ICF | independent colony founding | Wheeler 1913; Creighton 1950; JT Longino and PS Ward unpubl. collection data |
| F0035 | Formica mesasiatica | exsecta | free-living, TSP | unknown | monodomous, polydomous | TSP, DCF: budding | facultative temporary parasitism, high frequency budding | Seifert 2000 |
| F0129 | Formica moki | "fusca" | free-living | monogynous, polygynous | monodomous | ICF | independent colony founding | Francoeur 1973; SP Cover unpubl. collection data |
| F0117 | Formica montana nr. | "fusca" | free-living | monogynous, polygynous | monodomous, polydomous | ICF: haplometrosis, pleometrosis, DCF: budding | independent colony founding | Francoeur 1973; SP Cover unpubl. collection data |
| F0106 | Formica neoclara | "fusca" | free-living | monogynous, polygynous | monodomous | ICF | independent colony founding | Francoeur 1973; SP Cover unpubl. collection data |
| F0038 | Formica neogagates | neogagates | free-living | monogynous, polygynous | monodomous | ICF: haplometrosis, claustral | independent colony founding | Creighton 1940; Frumhoff and Ward 1992; SP Cover unpubl. collection data |
| F0107 | Formica neorufibarbis | "fusca" | free-living | monogynous, polygynous | monodomous | ICF | independent colony founding | Francoeur 1973; SP Cover unpubl. collection data |
| F0109 | Formica neorufibarbis | "fusca" | free-living | monogynous, polygynous | monodomous | ICF | independent colony founding | Francoeur 1973; SP Cover unpubl. collection data |
| F0127 | Formica neorufibarbis | "fusca" | free-living | monogynous, polygynous | monodomous | ICF | independent colony founding | Francoeur 1973; SP Cover unpubl. collection data |
| F0128 | Formica neorufibarbis | "fusca" | free-living | monogynous, polygynous | monodomous | ICF | independent colony founding | Francoeur 1973; SP Cover unpubl. collection data |
| F0108 | Formica neorufibarbis | "fusca" | free-living | monogynous, polygynous | monodomous | ICF | independent colony founding | Francoeur 1973; SP Cover unpubl. collection data |
| F0126 | Formica neorufibarbis | "fusca" | free-living | monogynous, polygynous | monodomous | ICF | independent colony founding | Francoeur 1973; SP Cover unpubl. collection data |
| F0077 | Formica nepticula | difficilis | free-living, TSP | monogynous, polygynous | monodomous | TSP | facultative temporary parasitism, low frequency budding | Wheeler 1913; SP Cover unpubl. collection data |
| F0078 | Formica nevadensis | difficilis | free-living, TSP | monogynous, polygynous | monodomous | TSP | facultative temporary parasitism, low frequency budding | Wheeler 1904; SP Cover unpubl. collection data |
| F0079 | Formica nevadensis | difficilis | free-living, TSP | monogynous, polygynous | monodomous | TSP | facultative temporary parasitism, low frequency budding | Wheeler 1904; SP Cover unpubl. collection data |
| F0080 | Formica nevadensis | difficilis | free-living, TSP | monogynous, polygynous | monodomous | TSP | facultative temporary parasitism, low frequency budding | Wheeler 1904; SP Cover unpubl. collection data |
| F0007 | Formica obscuripes | integra | free-living, TSP | monogynous, polygynous | monodomous, polydomous, supercolonial | TSP, DCF: budding | facultative temporary parasitism, high frequency budding | Frumhoff and Ward 1992; Talbot 2012 |
| F0008 | Formica obscuripes | integra | free-living, TSP | monogynous, polygynous | monodomous, polydomous, supercolonial | TSP, DCF: budding | facultative temporary parasitism, high frequency budding | Frumhoff and Ward 1992; Talbot 2012 |
| F0009 | Formica obscuripes | integra | free-living, TSP | monogynous, polygynous | monodomous, polydomous, supercolonial | TSP, DCF: budding | facultative temporary parasitism, high frequency budding | Frumhoff and Ward 1992; Talbot 2012 |
| F0162 | Formica obscuripes | integra | free-living, TSP | monogynous, polygynous | monodomous, polydomous, supercolonial | TSP, DCF: budding | facultative temporary parasitism, high frequency budding | Frumhoff and Ward 1992; Talbot 2012 |
| F0006 | Formica obscuriventris | integra | free-living, TSP | monogynous, polygynous | monodomous, polydomous | TSP | facultative temporary parasitism, low frequency budding | Creighton 1940; Frumhoff and Ward 1992; SP Cover and C Rabeling unpubl. collection data |
| F0011 | Formica obscuriventris | integra | free-living, TSP | monogynous, polygynous | monodomous, polydomous | TSP | facultative temporary parasitism, low frequency budding | Creighton 1940; Frumhoff and Ward 1992; SP Cover and C Rabeling unpubl. collection data |
| F0046 | Formica obtusopilosa | neogagates | free-living | monogynous, polygynous | monodomous | ICF: haplometrosis, claustral | independent colony founding | Creighton 1950; SP Cover unpubl. collection data |
| F0047 | Formica obtusopilosa | neogagates | free-living | monogynous, polygynous | monodomous | ICF: haplometrosis, claustral | independent colony founding | Creighton 1950; SP Cover unpubl. collection data |
| F0131 | Formica occulta nr. | "fusca" | free-living | monogynous, polygynous | monodomous, polydomous | ICF: haplometrosis | independent colony founding | Francoeur 1973; Frumhoff and Ward 1992; SP Cover unpubl. collection data |
| F0134 | Formica orangea | "fusca" | free-living | unknown | unknown | unknown | unknown | Seifert and Schultz 2009 |
| F0012 | Formica oreas | integra | free-living, TSP | monogynous, polygynous | monodomous, polydomous | TSP | facultative temporary parasitism, low frequency budding | Sather 1972; SP Cover and ML Borowiec unpubl. collection data |
| F0013 | Formica oreas | integra | free-living, TSP | monogynous, polygynous | monodomous, polydomous | TSP | facultative temporary parasitism, low frequency budding | Sather 1972; SP Cover and ML Borowiec unpubl. collection data |
| F0125 | Formica oreas | integra | free-living, TSP | monogynous, polygynous | monodomous, polydomous | TSP | facultative temporary parasitism, low frequency budding | Sather 1972; SP Cover and ML Borowiec unpubl. collection data |
| F0135 | Formica pacifica | "fusca" | free-living | unknown | unknown | unknown | unknown | Francoeur 1973; JT Longino unpubl. collection data |
| F0060 | Formica pallidefulva | pallidevulva | free-living | monogynous, polygynous | monodomous | ICF: haplometrosis, pleometrosis | independent colony founding | Trager et al. 2007; SP Cover unpubl. collection data |
| F0066 | Formica pallidefulva | pallidevulva | free-living | monogynous, polygynous | monodomous | ICF: haplometrosis, pleometrosis | independent colony founding | Trager et al. 2007; SP Cover unpubl. collection data |
| F0053 | Formica pergandei | sanguinea | TSP, obl. dulosis | monogynous, polygynous | monodomous | TSP | obligate temporary parasitism without budding (dulosis) | Trager et al. 2007; SP Cover unpubl. collection data |
| F0051 | Formica nr pergandei | sanguinea | TSP, obl. dulosis | monogynous, polygynous | monodomous | TSP | obligate temporary parasitism without budding (dulosis) | Trager et al. 2007; SP Cover unpubl. collection data |
| F0039 | Formica perpilosa | neogagates | free-living | monogynous, polygynous | monodomous | ICF: haplometrosis, claustral | independent colony founding | Wheeler 1913; SP Cover and C Rabeling unpubl. collection data |
| F0028 | Formica cf planipilis | integra | free-living, TSP | monogynous, polygynous | monodomous, polydomous, supercolonial | TSP, DCF: budding | facultative temporary parasitism, high frequency budding | Creighton 1940; SP Cover unpubl. collection data |
| F0145 | Formica podzolica | "fusca" | free-living | monogynous, polygynous | monodomous, polydomous | ICF: haplometrosis, DCF: budding | independent colony founding | Francoeur 1973; Frumhoff and Ward 1992; SP Cover unpubl. collection data |
| F0019 | Formica polyctena | rufa | free-living, TSP | monogynous, polygynous | monodomous, polydomous, supercolonial | TSP, DCF: budding | facultative temporary parasitism, high frequency budding | Collingwood 1979; Ito and Higashi 1990; Seifert 2018 |
| F0020 | Formica pratensis | rufa | free-living, TSP | monogynous, polygynous | monodomous, polydomous, supercolonial | TSP, DCF: budding | facultative temporary parasitism, high frequency budding | Horstmann et al. 1994; Sorensen and Schmidt 1983; Seifert 2018 |
| F0033 | Formica pressilabris | exsecta | free-living, TSP | monogynous, polygynous | monodomous, polydomous | TSP, DCF: budding | facultative temporary parasitism, high frequency budding | Pamilo and Rosengren 1984; Seifert 2018 |
| F0116 | Formica propatula | "fusca" | free-living | unknown | unknown | unknown | unknown | Francoeur 1973 |
| F0016 | Formica propinqua | integra | free-living, TSP | monogynous, polygynous | monodomous | TSP | facultative temporary parasitism, low frequency budding | Creighton 1940; SP Cover unpubl. collection data |
| F0056 | Formica puberula | sanguinea | TSP, obl. dulosis | monogynous, polygynous | monodomous | TSP | obligate temporary parasitism without budding (dulosis) | Creighton 1950; SP Cover unpubl. collection data |
| F0122 | Formica puberula | sanguinea | TSP, obl. dulosis | monogynous, polygynous | monodomous | TSP | obligate temporary parasitism without budding (dulosis) | Creighton 1950; SP Cover unpubl. collection data |
| F0081 | Formica querquetulana | difficilis | free-living, TSP | monogynous, polygynous | monodomous | TSP | facultative temporary parasitism, low frequency budding | Kennedy and Dennis 1937; SP Cover unpubl. collection data |
| F0095 | Formica querquetulana | difficilis | free-living, TSP | monogynous, polygynous | monodomous | TSP | facultative temporary parasitism, low frequency budding | Kennedy and Dennis 1937; SP Cover unpubl. collection data |
| F0026 | Formica ravida | integra | free-living, TSP | monogynous, polygynous | monodomous, polydomous | TSP, DCF: budding | facultative temporary parasitism, high frequency budding | Creighton 1940; SP Cover unpubl. collection data |
| F0027 | Formica ravida | integra | free-living, TSP | monogynous, polygynous | monodomous, polydomous | TSP, DCF: budding | facultative temporary parasitism, high frequency budding | Creighton 1940; SP Cover unpubl. collection data |
| F0086 | Formica reflexa | dakotensis | TSP with obligate dependence on host workers; possible inquilinism. Only mixed colonies; no evidence of dulosis | monogynous | monodomous | TSP, perhaps obligatory | permanent social parasitism | Buren 1942, SP Cover unpubl. collection data |
| F0054 | Formica rubicunda | sanguinea | free-living, TSP, obl. dulosis | monogynous, polygynous | monodomous | TSP | obligate temporary parasitism without budding (dulosis) | Creighton 1950; SP Cover unpubl. collection data |
| F0055 | Formica rubicunda | sanguinea | free-living, TSP, obl. dulosis | monogynous, polygynous | monodomous | TSP | obligate temporary parasitism without budding (dulosis) | Creighton 1950; SP Cover unpubl. collection data |
| F0018 | Formica rufa | rufa | free-living, TSP | monogynous, polygynous | monodomous, polydomous, supercolonial | TSP, DCF: budding | facultative temporary parasitism, high frequency budding | Collingwood 1979; Ito and Higashi 1990; Gösswald 2012; Seifert 2018 |
| F0136 | Formica rufibarbis | "fusca" | free-living | monogynous, polygynous | monodomous | ICF: haplometrosis, pleometrosis | independent colony founding | Gammans 2008; Seifert and Schultz 2009 |
| F0048 | Formica sanguinea | sanguinea | free-living, TSP, fac. dulosis | monogynous, polygynous | monodomous, polydomous | TSP, DCF: budding | obligate temporary parasitism without budding (dulosis) | Collingwood 1979; Frumhoff and Ward 1992; Mori and Le Moli 1998; Seifert 2018 |
| F0049 | Formica sanguinea | sanguinea | free-living, TSP, fac. dulosis | monogynous, polygynous | monodomous, polydomous | TSP, DCF: budding | obligate temporary parasitism without budding (dulosis) | Collingwood 1979; Frumhoff and Ward 1992; Mori and Le Moli 1998; Seifert 2018 |
| F0132 | Formica sibylla | "fusca" | free-living | unknown | monodomous | unknown | unknown | Francoeur 1973; PS Ward unpubl. collection data |
| F0133 | Formica cf sibylla | "fusca" | free-living | unknown | unknown | unknown | unknown | PS Ward unpubl. collection data |
| F0101 | Formica sp. "fusca grp" | "fusca" | free-living | unknown | unknown | unknown | unknown | SP Cover unpubl. collection data |
| F0142 | Formica sp. "fusca grp" | "fusca" | free-living | unknown | unknown | unknown | unknown | SP Cover unpubl. collection data |
| F0143 | Formica sp. "fusca grp" | "fusca" | free-living | unknown | unknown | unknown | unknown | SP Cover unpubl. collection data |
| F0160 | Formica sp. "fusca grp" | "fusca" | free-living | unknown | unknown | unknown | unknown | SP Cover unpubl. collection data |
| F0161 | Formica sp. "fusca grp" | "fusca" | free-living | unknown | unknown | unknown | unknown | SP Cover unpubl. collection data |
| F0165 | Formica sp. "fusca grp" | "fusca" | free-living | unknown | unknown | unknown | unknown | SP Cover unpubl. collection data |
| F0074 | Formica sp. "micro grp" | difficilis | free-living, TSP | monogynous, polygynous | monodomous | TSP | facultative temporary parasitism, low frequency budding | SP Cover unpubl. collection data |
| F0087 | Formica sp. "micro grp" | difficilis | free-living, TSP | monogynous, polygynous | monodomous | TSP | facultative temporary parasitism, low frequency budding | SP Cover unpubl. collection data |
| F0089 | Formica sp. "micro grp" | difficilis | free-living, TSP | monogynous, polygynous | monodomous | TSP | facultative temporary parasitism, low frequency budding | SP Cover unpubl. collection data |
| F0090 | Formica sp. "micro grp" | difficilis | free-living, TSP | monogynous, polygynous | monodomous | TSP | facultative temporary parasitism, low frequency budding | SP Cover unpubl. collection data |
| F0091 | Formica sp. "micro grp" | difficilis | free-living, TSP | monogynous, polygynous | monodomous | TSP | facultative temporary parasitism, low frequency budding | SP Cover unpubl. collection data |
| F0092 | Formica sp. "micro grp" | difficilis | free-living, TSP | monogynous, polygynous | monodomous | TSP | facultative temporary parasitism, low frequency budding | SP Cover unpubl. collection data |
| F0093 | Formica sp. "micro grp" | difficilis | free-living, TSP | monogynous, polygynous | monodomous | TSP | facultative temporary parasitism, low frequency budding | SP Cover unpubl. collection data |
| F0094 | Formica sp. "micro grp" | difficilis | free-living, TSP | monogynous, polygynous | monodomous | TSP | facultative temporary parasitism, low frequency budding | SP Cover unpubl. collection data |
| F0096 | Formica sp. "micro grp" | difficilis | free-living, TSP | monogynous, polygynous | monodomous | TSP | facultative temporary parasitism, low frequency budding | SP Cover unpubl. collection data |
| F0157 | Formica sp. "micro grp" | difficilis | free-living, TSP | monogynous, polygynous | monodomous | TSP | facultative temporary parasitism, low frequency budding | SP Cover unpubl. collection data |
| F0058 | Formica sp. "neogagates grp" | neogagates | free-living | monogynous, polygynous | monodomous | ICF: haplometrosis, claustral | independent colony founding | SP Cover unpubl. collection data |
| F0082 | Formica sp. mlb01 "micro grp" | difficilis | free-living, TSP | monogynous, polygynous | monodomous | TSP | facultative temporary parasitism, low frequency budding | ML Borowiec unpubl. collection data |
| F0083 | Formica sp. mlb02 "micro grp" | difficilis | free-living, TSP | monogynous, polygynous | monodomous | TSP | facultative temporary parasitism, low frequency budding | ML Borowiec unpubl. collection data |
| F0084 | Formica sp. mlb03 "micro grp" | difficilis | free-living, TSP | monogynous, polygynous | monodomous | TSP | facultative temporary parasitism, low frequency budding | ML Borowiec unpubl. collection data |
| F0085 | Formica sp. mlb04 "micro grp" | difficilis | free-living, TSP | monogynous, polygynous | monodomous | TSP | facultative temporary parasitism, low frequency budding | ML Borowiec unpubl. collection data |
| F0123 | Formica sp. sanguinea grp | sanguinea | TSP, dulosis (fac. or obl.?) | unknown | unknown | unknown | unknown | SP Cover unpubl. collection data |
| F0104 | Formica subaenescens | "fusca" | free-living | monogynous, polygynous | monodomous, polydomous | ICF: haplometrosis, pleometrosis | independent colony founding | Francoeur 1973; SP Cover unpubl. collection data |
| F0022 | Formica subnitens | integra | free-living, TSP | monogynous, polygynous | monodomous, polydomous | TSP | facultative temporary parasitism, low frequency budding | Creighton 1940; SP Cover unpubl. collection data |
| F0025 | Formica subnitens | integra | free-living, TSP | monogynous, polygynous | monodomous, polydomous | TSP | facultative temporary parasitism, low frequency budding | Creighton 1940; SP Cover unpubl. collection data |
| F0014 | Formica cf subnitens | integra | free-living, TSP | monogynous, polygynous | monodomous, polydomous | TSP | facultative temporary parasitism, low frequency budding | Creighton 1940; SP Cover unpubl. collection data |
| F0137 | Formica subpolita | "fusca" | free-living | monogynous, polygynous | monodomous, polydomous | ICF: haplometrosis | independent colony founding | Francoeur 1973; SP Cover unpubl. collection data |
| F0138 | Formica subpolita | "fusca" | free-living | monogynous, polygynous | monodomous, polydomous | ICF: haplometrosis | independent colony founding | Francoeur 1973; SP Cover unpubl. collection data |
| F0144 | Formica subsericea | "fusca" | free-living | monogynous, polygynous | monodomous, polydomous | ICF: haplometrosis, pleometrosis | independent colony founding | Francoeur 1973; Frumhoff and Ward 1992; SP Cover unpubl. collection data |
| F0146 | Formica subsericea | "fusca" | free-living | monogynous, polygynous | monodomous, polydomous | ICF: haplometrosis, pleometrosis | independent colony founding | Francoeur 1973; Frumhoff and Ward 1992; SP Cover unpubl. collection data |
| F0159 | Formica subsericea | "fusca" | free-living | monogynous, polygynous | monodomous, polydomous | ICF: haplometrosis, pleometrosis | independent colony founding | Francoeur 1973; Frumhoff and Ward 1992; SP Cover unpubl. collection data |
| F0156 | Formica talbotae | difficilis | workerless inquilinism | polygynous | monodomous | TSP | permanent social parasitism | Talbot 1977, Wilson 1977, Talbot 2012 |
| F0141 | Formica transmontanis | "fusca" | free-living | unknown | monodomous | unknown | independent colony founding | Francoeur 1973; PS Ward unpubl. collection data |
| F0023 | Formica truncorum | rufa | free-living, TSP | monogynous, polygynous | monodomous, polydomous, supercolonial | TSP, DCF: budding | facultative temporary parasitism, high frequency budding | Collingwood 1979; Frumhoff and Ward, 1992; Seifert 2018 |
| F0029 | Formica ulkei | exsecta | free-living, TSP | monogynous, polygynous | monodomous, polydomous, supercolonial | TSP, DCF: budding | facultative temporary parasitism, high frequency budding | Holmquist 1928; SP Cover unpubl. collection data |
| F0024 | Formica uralensis | uralensis | free-living, TSP | monogynous, polygynous | monodomous, polydomous, supercolonial | TSP, DCF: budding | facultative temporary parasitism, high frequency budding | Busch 1998; Münch 1991; Preuß 1956; Preuß 1957; Seifert 2018 |
| F0057 | Formica wheeleri | sanguinea | TSP, obl. dulosis | monogynous, polygynous | monodomous | TSP | obligate temporary parasitism without budding (dulosis) | Creighton 1950; SP Cover unpubl. collection data |
| F0139 | Formica xerophila | "fusca" | free-living | monogynous | monodomous | ICF: haplometrosis | independent colony founding | Francoeur 1973; SP Cover unpubl. collection data |
| F0140 | Formica cf xerophila | "fusca" | free-living | monogynous | monodomous | ICF: haplometrosis | independent colony founding | PS Ward unpubl. collection data |
| D1135 | Iberoformica subrufa | n/a | free-living | monogynous | monodomous | ICF | independent colony founding | Tinaut 1990; Muñoz-López et al. 2012; Seifert 2018 |
| F0158 | Polyergus longicornis | n/a | TSP, obl. dulosis | monogynous | monodomous | TSP | obligate temporary parasitism without budding (dulosis) | Trager 2013 |
| F0153 | Polyergus mexicanus | n/a | TSP, obl. dulosis | monogynous | monodomous | TSP | obligate temporary parasitism without budding (dulosis) | Trager 2013 |
| F0154 | Polyergus mexicanus | n/a | TSP, obl. dulosis | monogynous | monodomous | TSP | obligate temporary parasitism without budding (dulosis) | Trager 2013 |
| F0149 | Polyergus rufescens | n/a | TSP, obl. dulosis | monogynous | monodomous | TSP | obligate temporary parasitism without budding (dulosis) | Trager 2013 |
| F0124 | Polyergus vinosus | n/a | TSP, obl. dulosis | monogynous | monodomous | TSP | obligate temporary parasitism without budding (dulosis) | Trager 2013 |
| F0150 | Proformica mongolica | n/a | free-living | monogynous, polygynous | monodomous | ICF | independent colony founding | Galkowski et al. 2017 |
| F0152 | Rossomyrmex minuchae | n/a | TSP, obl. dulosis | monogynous | monodomous | TSP | obligate temporary parasitism without budding (dulosis) | Ruano and Tinaut 2005 |
